# DNA Nanocage-Mediated Aspirin Delivery Mitigates Excitotoxicity and Promotes Neuroprotection in Ischemic Stroke-Induced Zebrafish

**DOI:** 10.64898/2025.12.23.696221

**Authors:** Krupa Kansara, Shaivee Chokshi, Keya Jantrania, Sandip Mandal, Prabal Kumar Maiti, Ashutosh Kumar, Dhiraj Bhatia

**Affiliations:** Department of Biosciences and Bioengineering, Indian Institute of Technology Gandhinagar, Gujarat 382355, India; Biological and Life Sciences, School of Arts and Sciences, Ahmedabad University, Ahmedabad, Gujrat 380009, India; Centre for Condensed Matter Theory, Department of Physics, Indian Institute of Science, Bangalore, India

**Author notes:** Contributed equally (Shaivee Chokshi, Keya Jantrania, Sandip Mandal).

**Keywords:** DNA tetrahedron, Aspirin, MD Simulation, zebrafish, ischemic stroke, neuroprotection, apoptosis, ROS, nanotherapeutics

## Abstract

Oxidative stress, apoptosis, excitotoxicity, and compromised neuronal function are the hallmarks of ischaemic stroke, a major cause of neurological impairment. Because of their controlled-release properties and biocompatibility, DNA tetrahedron nanostructures (TD) offer a viable platform for targeted drug delivery. In this work, we used a zebrafish larval model of ischaemic stroke to assess the neuroprotective potential of TD-mediated aspirin administration and investigated the TD:Aspirin complexation modes using molecular dynamics (MD) simulations. In addition to morphological abnormalities, elevated reactive oxygen species (ROS), thrombus formation, intracellular calcium accumulation, and dysregulated expression of neuroprotective and apoptotic genes, stroke induction resulted in significant behavioural deficits, including decreased locomotor activity, increased latency, and erratic swimming. Treatment with TD:Aspirin, especially at a 1:100 ratio, markedly improved locomotor behaviour, restored production of BDNF, GDNF, MBP, α-tubulin, GRIN2B, SLC8A, NOS1, and caspases, normalised heart rate, lowered ROS generation and apoptosis, and decreased thrombus and calcium deposition. TD:Aspirin showed better efficacy than free Aspirin and TD-only groups, perhaps as a result of improved stability, bioavailability, and sustained release. These results demonstrate the potential of drug delivery via DNA nanocages as a novel therapeutic approach for ischaemic stroke, offering neuroprotection, functional recovery, and neuronal homeostasis restoration.

**TOC Graphic:** DNA Nanocage-Mediated Aspirin Delivery Mitigates Excitotoxicity and Promotes Neuroprotection in Ischemic Stroke-Induced Zebrafish

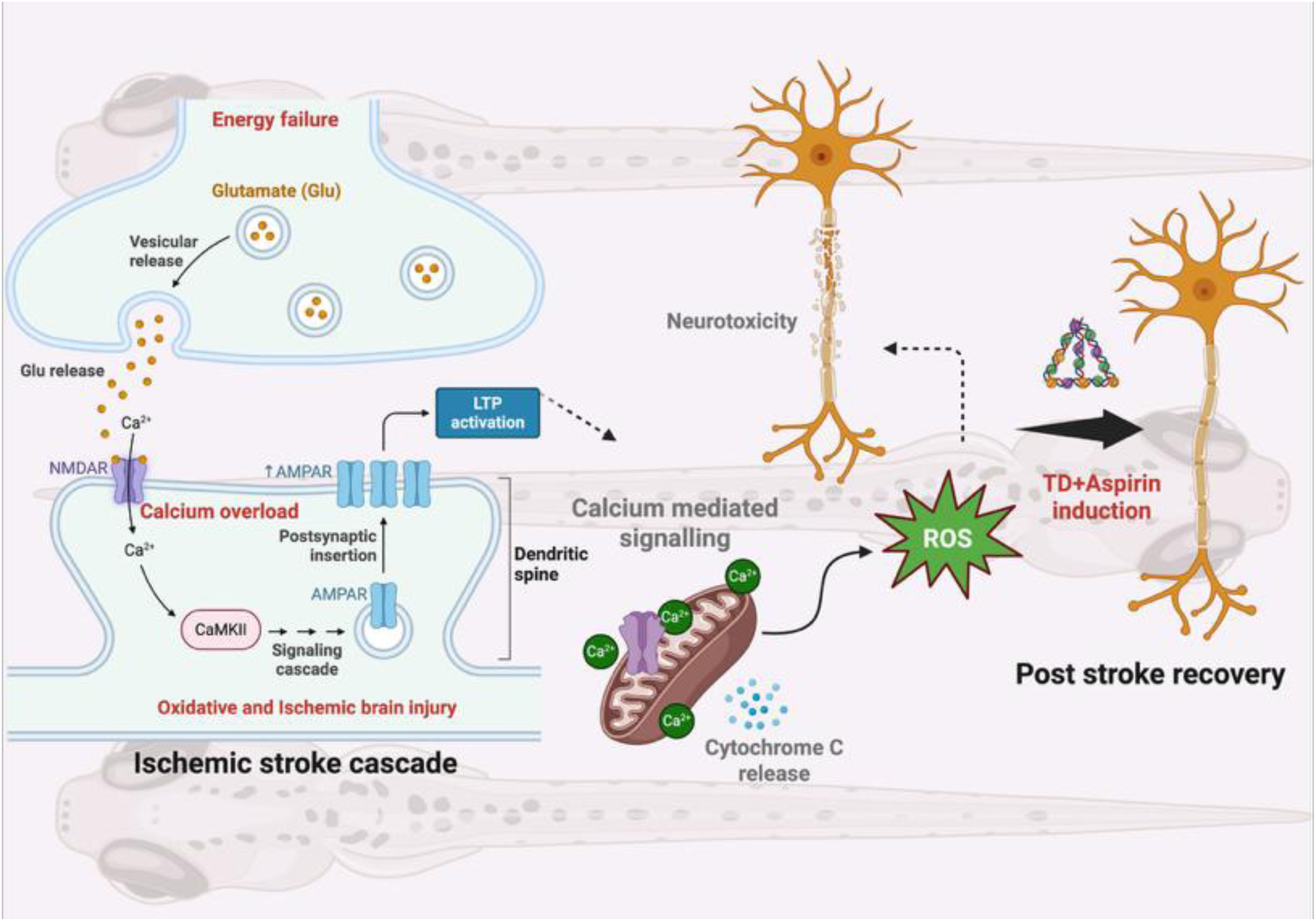

## 1. Introduction

Clinically, a stroke can be defined as a sudden onset of focal neurological dysfunction of the central nervous system^1, 2^. Ischaemic stroke continues to be one of the world’s leading causes of death and permanent neurological impairment^3^. The development of ischemic stroke (IS) is contributed to several risk factors, such as diabetes, smoking, high cholesterol levels, and high blood pressure. Based on the underlying cause, the resulting IS may result from an embolism which can either originate in the heart or travel from one artery to another, or due to a disease which affects the small blood vessels within the brain^4^. Antithrombotic agents like clopidogrel, ticlopidine, extended-release dipyridamole, warfarin, and aspirin have proven to be effective in the prevention of ischemic stroke^5^. However, tissue plasminogen activator (tPA) is the only FDA-approved drug used for the treatment of acute thrombotic stroke. This therapy, according to the NIH, is currently received only by 1-3% of patients in the US^6^. Moreover, there are only a limited number of stroke patients who are qualified for the current therapy because of the narrow therapeutic windows, co-morbidities, and limited hospital resources. Ischemic stroke patients can get the tPA therapy within 4.5 hours of symptom onset, however, only 10-15% patients are eligible. Patients with proximal large vessel occlusion, can undergo surgical thrombectomy within 6 to 24 hours^7^. The therapeutic choices are still restricted because of the short treatment windows, systemic adverse effects, and poor penetration of medications across the blood–brain barrier (BBB), even with the availability of pharmacological therapies such as antiplatelet and thrombolytic agents. In order to effectively manage stroke, there is a rising demand for innovative nanocarrier systems that can enhance the therapeutic drugs’ bioavailability, target specificity, and controlled release.

Among antithrombotic agents, aspirin is popularly prescribed since it needs less monitoring, is easily available, and is low-cost. Aspirin is often prescribed initially, and according to literature, it is the only antiplatelet drug that is beneficial in acute ischemic stroke ^8^. The Chinese Acute Stroke Trial (CAST) and the International Stroke Trial (IST) are two major trials that provide evidence supporting the efficacy of aspirin in ischemic stroke^5^. The findings from these trials suggest that aspirin, when consumed between 160-300 mg within 48 hrs of ischemic stroke, provides small but certain reductions in death and recurrent ischemic stroke rates. The American College of Chest Physicians (ACCP) has recommended early aspirin therapy - Grade 1A. This therapy includes giving aspirin at doses of 150-325 mg for patients who have had an acute ischemic stroke and have not received thrombolysis with tissue plasminogen activation. The American Stroke Association (ASA) and the American Heart Association (AHA) guidelines for early management of acute ischemic stroke recommend that the initial dose of aspirin be 325 mg. Aspirin doses from the range of 30 to 1300 mg daily have proven to prevent strokes in patients with transient ischemic stroke (TIA) or stroke; higher doses however are linked with risk of gastrointestinal hemorrhage. Minimum dosage for secondary stroke prevention is 50 mg daily according to ESPS-2 trials, while AHA/ASA guidelines recommend 50 - 325 mg daily for the same^9, 10^.

Several intricate ischaemic cascades, such as the release of excitatory neurotransmitters, oxidative stress damage, and inflammatory responses, take place in the local brain tissue during ischaemia and reperfusion^11, 12^. The metabolism of proteins, lipids, and nucleic acids is disrupted by excitotoxicity and oxidative stress, which generate an intracellular calcium overload and a significant amount of reactive oxygen species (ROS) in neurons ^13–15^. Furthermore, the ischaemic hemisphere’s local microenvironment deteriorates due to the release of pro-inflammatory molecules such TNF-α and IL-1β, which exacerbates neurological dysfunction and neuronal death. Consequently, neuroprotective medicines that increase neuronal resistance to ischaemia and extend the window for intervention in the ischaemic cascades, when used in conjunction with recanalised therapy, can reduce neuronal mortality in the penumbra and are prospective treatment approaches for IS. For brain damage, nanomaterials have been employed as neuroprotective agents. However, the therapeutic use of nanotechnology in the treatment of brain injury is still in its infancy because of the complexity of the nervous system and the neurotoxicity of materials. According to previous studies, the DNA tetrahedrons (TDs) play active biological roles in neuroprotection and as an anti-inflammatory agent due to their easy synthesis and good biocompatibility, which is determined by their DNA composition ^16^. Owing to the advantages of TDs, they have been widely used in the field of biomedicine^17, 18^, especially in neuro-medicine^19, 20^. Several studies have observed that the TDs are able to promote the proliferation, migration, neuronal differentiation of neural stem cells (NSCs)^21^, and also showcases autophagy modulation in a precise manner^22^. It was also observed that synergistic therapy with TDs and NSCs has improved spinal cord injury recovery has been improved^23^. Since they are able to cross the blood-brain barrier (BBB), these nanocages are highly useful in treating neurological disorders. For the drug to transport into the brain tissue, it is essential to overcome the blood-brain barrier, and TDs have proven valuable for the delivery of various bioactive molecules^24^. In a study done by Yang and her colleagues on normal nude mice, they were able to demonstrate that, to some extent, TDs could cross the BBB^25^. Studies have also been carried out on zebrafish embryos, proving uptake of TDs ^19, 20, 26–28^.

In this study, four single-stranded oligonucleotides self-hybridised to produce a compact three-dimensional tetrahedral structure, which was used to design and build TDs. To the best of our knowledge, Aspirin (acetylsalicylic acid), a clinically proven antiplatelet and anti-inflammatory medication, was conjugated to the TDs framework to create TD: Aspirin nanoconjugates for the very first time, which increased therapeutic efficacy against ischaemic stroke as a novel nanoconjugate. Enhancing aspirin’s stability in physiological settings, improving its pharmacokinetic profile, lowering dosages and enabling targeted distribution across biological barriers were the goals. A zebrafish (*Danio rerio*) model of ischaemic stroke was used to assess these nanostructures’ neuroprotective potential. Because of their remarkable genetic and physiological similarity to humans, optical transparency during early development, and suitability for real-time imaging and behavioural analysis, zebrafish have become a reliable vertebrate model for researching cerebrovascular diseases. TDs and TD:Aspirin conjugates were used to treat the stroke-like disease created by chemically inducing ischaemic circumstances. Morphological observations, behavioural recovery analysis, and first neuroprotective evaluations were all part of the post-treatment evaluations. The physicochemical characterisation of the synthesised TDs and TD:Aspirin nanoconjugates, including electrophoretic mobility, size distribution, zeta potential, and topographical morphology, is documented in the dataset that goes with this publication. Additionally, we have used molecular dynamics (MD) simulations to investigate the mode of TD:Aspirin complexation mechanism and the structural stability of the TD nanocage upon Aspirin binding. It also records the detailed biological response profiles and excitotoxicity pathway in ischaemic stroke-induced zebrafish, providing important baseline information for upcoming translational research investigating medication delivery systems based on DNA nanostructures for cerebrovascular disorders.

## 2. Experimental section

### 2.1. Revealing the Distinctive Features of TDs and Formulated TDs: Aspirin

#### 2.1.1. Synthesis of TDs and TD:Aspirin Conjugations

DNA TD nanostructures were synthesised using a previously described one-pot synthesis method ^29^. A thermal annealing procedure was performed in a PCR apparatus with cycling settings from 95 to 4 °C, after four complementary single-stranded DNA oligos (M1, M2, M3, and M4) were combined in equimolar concentration with 2 mM MgCl_2_. After 10 minutes of heating to 95 °C, the reaction mixture was progressively cooled to 4 °C. Oligonucleotides labelled with cyanine-5 (Cy5) were employed for imaging. To functionalise the TD: Aspirin conjugations, a combination of DNA TD and Aspirin in a 1:50 and 1:100 molar ratio was incubated for 5 hours.

#### 2.1.2. Characterisation and conjugation of TDs and TDs with Aspirin

##### DLS and Zeta Potential

A Malvern analytical Zetasizer Nano-ZS was used to perform the DLS for hydrodynamic size and zeta potential measurements for charge distribution on TDs and TD: Aspirin conjugations (1:50). Following a Gaussian fit, the data were plotted on GraphPad Prism software.

##### Atomic Force Microscopy

The self-assembled tetrahedral DNA nanocages (TDs) and TD: Aspirin conjugations (1:50) were examined for morphological characteristics and structural integrity using Atomic Force Microscopy (AFM). The samples were prepared by following established lab protocols. 5−10 μL aliquots of TD and TD: Aspirin (1:50) were spread over a freshly sliced mica substrate to record the AFM pictures in tapping mode with RFESPG-75 (antimony-doped Si) and SCANASYST-AIR (silicon nitride) probes in ambient circumstances. The imaging was performed with a Bruker NanoWizard Sense + Bio AFM installed at IIT Gandhinagar, Gujarat.

##### Electrophoretic Mobility Shift Assay

Native-PAGE was used in an electrophoretic mobility shift assay. To investigate the creation of the tetrahedral structure, a 10% polyacrylamide gel was made. After 90 minutes of operation at 90 V, the gel was stained with EtBr and examined using the Bio-Rad ChemiDoc MP imaging system for gel documentation.

### 2.2. Determination of Aspirin entrapment efficiency in TDs and kinetic release

The TD: Aspirin (1:50) were stored inside the dialysis membrane, and this assembly was then immersed in a beaker containing phosphate buffer with a pH of 5.5, 7.4 and 9.4 while being continuously stirred. After a 12-hour incubation period, the phosphate buffer solution was collected to determine the entrapment efficiency. Subsequently, the phosphate buffer was replaced with fresh phosphate buffer (pH 5.5, 7.4 and 9.4), and the setup was maintained for an additional 48 hours to collect hourly samples (1hr to 8hr, 12hr, 24hr, and 48hr) for drug release analysis. The drug content of the collected solutions was measured using a spectrophotometer. While the kinetic drug release profile was determined, the Entrapment Efficiency (%EE) was computed using the amount of drug released over the first 24 hours.

### 2.3. Molecular Dynamics Simulations

The initial atomistic structure of the tetrahedral DNA (TD) was constructed using the Polygen^30^, while the Aspirin molecule (PubChem CID: 2244) was obtained from the PubChem database. The geometry of the Aspirin molecule was optimized with the B3LYP functional together with a 6-311G* basis set using Gaussian 16 program. All the built systems were solvated in a cubic water box with a minimum padding of 15 Å using the TIP3P^31^ water model with xLEAP module of AMBER20^32^. As a result, all the solute atoms will be 15 Å away from the edge of the water box, ensuring adequate hydration. The negative charges of the DNA phosphate backbones were neutralized by adding the appropriate number of Mg^2+^ ions. To attain the desired salt concentration of 0.15 mM, we added appropriate numbers of MgCl_2_ molecules. Periodic boundary conditions were applied in all three directions to replicate the bulk properties of the system. All MD simulations were performed using PMEMD.CUDA module of AMBER20. The interactions of TD-DNA atoms were described by leaprc.DNA.bsc1 forcefield parameters ^33, 34^. For the divalent (Mg^2+^) ions, the Li-Merz ion parameters were applied. The interaction parameters for the non-standard Aspirin molecules were generated from the GAFF2 ^35^, with partial atomic charges derived using a RESP fitting scheme^36^.

The energy minimization of the solvated systems was first carried out using of steepest descent, followed by the conjugate gradient for the next 6000 steps. During this minimization stage, all TD and Aspirin atoms were restrained with a harmonic potential force constant of 500 Kcal.mol^-1^. Å^-2^, allowing the ions and water molecules to relax and remove any unfavourable contacts. The positional restraints on TD and Aspirin atoms were then gradually reduced from 500 to 5 kcal.mol^-1^.Å^-2^ in five successive steps, leading to full equilibration without constraints.

The systems were subsequently heated slowly in four steps: first, from 10 to 50 K for 6,000 MD steps, secondly, from 50 to 100 K for 12,000 steps, from 100 to 200 K for 10,000 steps, from 200 to 300 K for 10,000 steps, and finally, held at 300 K for another 12000 MD steps. Position restraints were applied to all the heavy atoms of the solute during heating with a force constant of 20 kcal.mol^-1^. Å ^-2^. Following heating process, the systems undergo a 5 ns of NPT equilibration using Berendsen weak coupling method at a target pressure of 1 bar^37, 38^. All the hydrogen-containing bonds were constrained using the Shake algorithm, which allowing us to use an integration time-step of 2 fs. Long-range electrostatic interactions were treated using the Particle Mesh Ewald (PME) method with a cutoff distance of 10 Å^39^. Following the equilibration, the systems were simulated for production run using a Langevin thermostat with a 1 ps of coupling constant for 200 ns at 300 K ^40^. Similar simulation protocols have been used in our earlier studies^41–46^. MD trajectory analysis was conducted using VMD and CPPTRAJ, while visualization and figure preparation were performed using XMGrace, PyMOL, and Python Matplotlib^47–49^.

### 2.4. Zebrafish maintenance

The laboratory environment was maintained at a temperature of 27°C-28°C, with a 14-hour light/10-hour dark cycle. Adult zebrafish were kept in 20-litre aquarium tanks with mildly saline water (Red Sea Coral Pro salt) and aeration provided by pumps. Using a pH meter (Horiba), the pH of the water was constantly maintained between 6.8 and 7.2. The fish were fed brine prawns (Artemia) twice daily and basic flakes (Aquafin) once a day. Breeding took place in a lab-built breeding box, with a male-to-female ratio of 3:4. To maximise egg production, fish were separated for at least two days before breeding twice weekly. Spawning happened within one hour of the divider being removed. After egg collection, they were kept in E3 medium, comprising 5.0 mM NaCl, 0.17 mM KCl, 0.33 mM CaCl₂, and 0.33 mM MgSO₄, and placed in sterile petri plates. These plates were kept in an incubator with temperature maintained at 28°C, until further experimentation. The experiments were conducted on 3-4 days post-fertilisation (dpf) larvae^50^.

### 2.5. Stroke generation via chemical hypoxia in zebrafish larvae

Healthy larvae of 3-4 dpf were selected for the experiments. Two groups of larvae were introduced to CoCl_2_ treatment: 10mM and 20mM. The concentration of CoCl_2_ was fixed based on a literature review, and experiments were conducted at 5mM, 10mM, 12mM, 15mM and 20mM ^51^. These larvae were closely monitored for behavioural and morphological changes. Several factors, such as increased latency, erratic swimming and reduced blood flow, were monitored to visually know if the stroke had been induced. The time frame for inducing hypoxia was fixed at 2 hours to ensure minimal mortality. Once induced, the larvae were introduced to three recovery treatments: 12.5μ M aspirin solution made in nuclease-free water, aspirin conjugated with DNA nanocages and E3 media^3^. The larvae were monitored for 3 hours posttreatment. Selected healthy larvae from the batch were used as a control group.

#### 2.5.1. Quantification of Ischemic stroke generation in zebrafish larvae Blood flow monitoring and Thrombus formation

O -Dianisidine staining was used to study the formation of thrombus and blood flow in the larvae. Treated larvae were stained following the protocol from Detrich et al ^52^(15 min in the dark in o-dianisidine (0.6 mg/ml), 0.01 M sodium acetate (pH 4.5),0.65% H_2_O_2_, and 40% (vol/vol) ethanol) with minor modifications. The larvae were stained with o-dianisidine, following a brief bleaching treatment using a solution of 0.8% KOH, 0,9% H_2_O_2_ and 0.1% Tween-20. The larvae were washed with PBST and stored in PFA. The imaging was done using a stereo microscope.

### 2.6. Morphological assessment of Ischemic stroke-induced zebrafish larvae

Ten to fifteen larvae at 3dpf were subjected to hypoxia treatment and later to recovery treatments: E3 media, 12.5μ g/ml aspirin, TD, TD: Aspirin (1:50, 1:100). Activity, latency, erratic swimming, mortality and malformations were observed and documented at three time stamps (1 hour, 1.5 hours and 2.5 hours) post treatment. Heart rate was measured 2 hours after hypoxia treatment and 2.5 hours after the recovery treatments. These changes were examined using a stereo microscope.

### 2.7. Intracellular reactive oxygen species evaluation in Ischemic stroke-induced zebrafish larvae

The oxidative stress in larvae was measured by quantifying the generation of reactive oxygen species (ROS). DCFDA dye was used to quantify ROS in the larvae. DCFDA dye was added at a concentration of 2mM in a 6-well plate. The plate was incubated for 1 hour at 28℃ in the dark. After incubation, three E3 media washes were given. Imaging was carried out in a fluorescent microscope equipped with a filter at 450nm - 490nm (blue light). The fluorescence intensities were measured using the ImageJ software^50^.

### 2.8. Apoptosis

Apoptotic cells in the larvae were visualised by staining with acridine orange dye. Larvae subjected to treatments were placed in a 6-well plate, and the acridine orange staining protocol was followed. The larvae were introduced to 10ug/ml of acridine orange stain in E3 media and the plate was incubated for 30 minutes at 28℃ in the dark. After incubation, the larvae were washed 3 times with E3 media. Imaging was carried out in a fluorescent microscope equipped with a filter at 450nm - 490nm. The fluorescence intensities were measured using the ImageJ software^50^.

### 2.9. Brain calcification assessment in Ischemic stroke-induced zebrafish larvae

The deposition of calcium in the neural crest due to ischemic stroke was quantified using the calcein dye that binds to calcium. The larvae were stained in calcein following the protocol by Zhao et al., with minor modifications^53^. Larvae subjected to treatments were washed with E3, and calcein prepared in MilliQ was added to the wells. The plate was incubated for 20 minutes in the dark at room temperature. The larvae were washed with E3 media 3 times before immediate imaging in a fluorescence microscope equipped with a 450nm-490nm filter.

### 2.10. Gene Expression Studies using qRT-PCR

Total RNA was isolated at two time stamps in the experiment. Initially, the RNA was isolated 2 hours post-hypoxia treatment, followed by RNA isolation for recovery treatment groups, 2.5 hours post-treatment. The process was carried out using a Favoregen RNA isolation kit. cDNA synthesis was carried out using 1000ng of isolated RNA, which was followed by a real-time PCR. The genes associated with ischemic stroke and its recovery were assessed such as BDNF (Brain-Derived Neurotrophic Factor), GDNF (Glial Cell Line-Derived Neurotrophic Factor), alpha-tubulin, and mbp (Myelin Basic Protein). Importantly, the genes associated with the excitotoxicity pathway were also analysed such as Grin2b (Glutamate Ionotropic Receptor NMDA Type Subunit 2B), SLc8a3 (Solute Carrier Family 8 Member A3), Nos1 (Nitric Oxide Synthase 1 (Neuronal NOS)), Dapk1(*Death-Associated Protein Kinase 1*), Caspase3 (*Cysteine-aspartic acid protease 3)*, Caspase9 (Cysteine-aspartic acid protease 9), and Bcl2 (B-cell Lymphoma 2). GAPDH was used as a housekeeping gene for both cases.

### 2.11. Statistical analysis

The experiments were carried out three times. The data were presented as the mean standard deviation for outcomes that were continuous. One-way and two-way analysis of variance (ANOVA) using GraphPad Prism was used to find statistically significant differences (P<0.05) between the three groups (version 5; La Jolla, CA, USA).

## 3. Results and Discussion

### 3.1. Characterisation and conjugation of TDs and TDs with Aspirin

The hydrodynamic size distribution and dispersity of the self-assembled TDs and TD: Aspirin (1:50) nanocages in solution were assessed using Dynamic Light Scattering (DLS). After appropriately diluting the samples in nuclease-free buffer to reduce multiple scattering effects, the measurements were carried out at 25 °C with a standard scattering angle of 90°. With an average hydrodynamic diameter of roughly 11.23 1.56 nm for TDs, 146.1 0.26 for TD: Asp (1:50) and 197.6 1.26 for TD: Asp (1:100), the DLS results showed a narrow and monodisperse size distribution that closely matches the predicted measurements of the intended TDs and TD: Asp conjugations. A high degree of homogeneity and effective self-assembly without notable aggregation or multimodal particle populations are suggested by the low polydispersity index (PDI < 0.25) (Figure 1B). The surface charge and colloidal stability of the TDs and TD: Asp (1:50, 1:100) were assessed using zeta potential analysis. The negatively charged phosphate backbone of DNA is compatible with the nanocages’ negative zeta potential, which was -36.32 ±1.22 mV. The successful conjugation of aspirin molecules on TDs showed the decreased zeta potential, such as -25.64 ±2.57 mV and - 20.46±1 for TD: Aspirin (1:50) and TD: Aspirin (1:100) (Figure 1C). In aqueous media, this negative potential helps to ensure colloidal stability and inhibit aggregation by promoting electrostatic repulsion between particles. The development of uniform, stable, and evenly distributed TDs and TD: Aspirin conjugation in solution are confirmed by the combined DLS and zeta potential measurements. Together with AFM observations, these results provide additional confirmation of the TDs’ effective synthesis and structural soundness.

**Figure 1:**
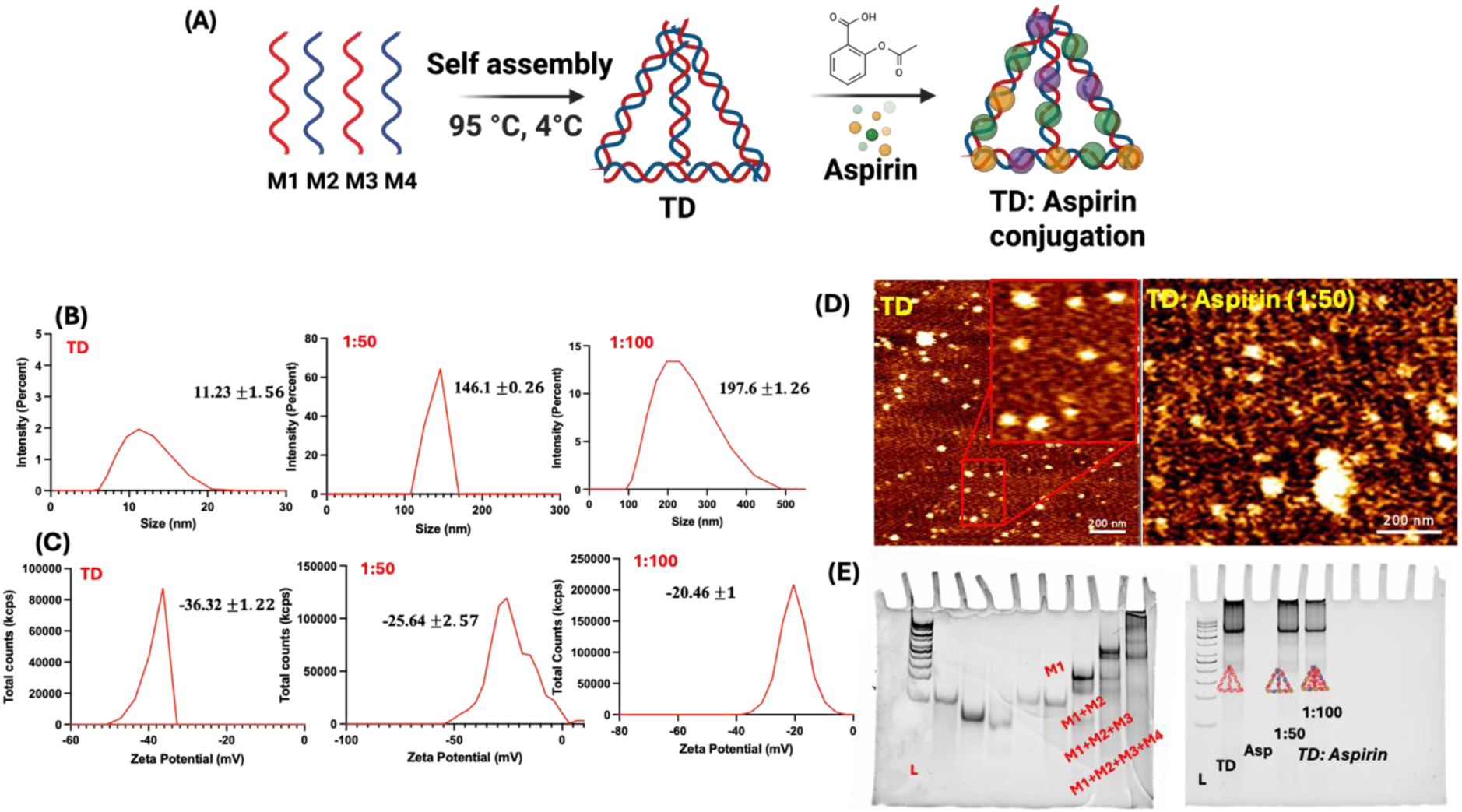
Characterization of a tetrahedral DNA nanostructure A) Schematic representation of the oligonucleotides self-assembly into Tetrahedron using the annealing process; B) DLS results of synthesized Tetrahedron showed the hydrodynamic diameter C) Zeta potential of negatively charged DNA nanoparticles D) AFM Images of DNA tetrahedron and TDN:Aspirin conjugation E) gel electrophoresis mobility shift-based characterization showing the retardation in the mobility upon formation of the Tetrahedron (T1+T2+T3+T4) and TDN:Aspirin (1:50), (1:100).

Uniformly dispersed nanostructures with clear, well-defined triangle or dot-like characteristics that corresponded to individual TDs and TD: Aspirin (1:50) were visible in AFM topography pictures. The measured lateral dimensions were in close agreement with the predicted size estimated from the specified DNA model, ranging from 12 to 15 nm. Partial flattening on the mica substrate and tip convolution effects are responsible for slight changes in the apparent size. The three-dimensional tetrahedral configuration was confirmed by height profile analysis, which showed an average height of 5-7 nm. High assembly fidelity and structural integrity are suggested by the discovered nanostructures’ homogeneity and monodispersity. The self-assembled TDs and TD: Aspirin (1:50) stability and durability under imaging circumstances were demonstrated by the lack of noticeable aggregation or deformation. The successful creation of distinct, uniform TDs and TD: Aspirin (1:50) nanocages with structural traits in line with the intended architecture was confirmed by AFM analysis. These results verify the accuracy of the DNA-based building at the nanoscale and support the effectiveness of the assembly procedure (Figure 1D).

The TDs and TD: Aspirin conjugations were successfully assembled in a stepwise manner, as shown by native polyacrylamide gel electrophoresis (PAGE) (Figute 1E). To confirm proper hybridisation and structure creation, the electrophoretic mobility of individual oligonucleotides, partial assemblies, and the completed TDs was compared. As the DNA strands gradually assembled, distinct mobility alterations were seen. Single-stranded oligonucleotides moved more quickly and formed bands close to the gel’s bottom. Intermediate bands representing partially formed structures emerged at higher places after complementary strands were sequentially mixed and annealed, suggesting a higher molecular weight and decreased electrophoretic mobility. The fully completed TDs sample showed a single, distinct band with the slowest migration, indicating that a compact, higher-order tetrahedral structure had successfully formed. However, the faint bands of TD: Asp (1:50, 1:100) were observed due to the aspirin conjugation over TDs. High assembly yield and structural consistency are suggested by the final lane’s lack of smeared or numerous bands. In contrast to single-stranded or partially assembled intermediates, the mobility pattern was in line with previously documented migratory behaviour of well-formed TDs, where compact, covalently connected junctions provide a noticeable retardation (Figure 1E).

### 3.2. TD-Aspirin Binding Modes from Molecular Dynamics

To complement experimental observation of TD:Aspirin complexation, we performed 200 ns atomistic MD simulations of TD in the presence of Aspirin molecules under physiological conditions for four systems: TD alone in bulk water, TD + 1 Aspirin, and TD + 100 and 150 Aspirin molecules (i.e., 1:100 and 1:150 ratios for TD:Aspirin) (Figure 2A-C). Our MD simulation results reveal the molecular origins of Aspirin interaction with TD. The TD nanocage is composed of 6 dsDNA helices and single unpaired nucleotides at the junction. Thus, TD has a structurally rich topology than the native dsDNA and provides more potential binding sites for Aspirin. For the single TD:Aspirin system, simulation results reveal that Aspirin molecule consistently binds to the outer corner mode of DNA-TD over the last ∼100 ns of the simulation time. This binding mode is driven by the π-π stacking interactions between unpaired TD nucleotides and aromatic rings of the Aspirin, highlighting strong binding affinity of the outer corner modes (Figure 2A). For the 1:100 TD:Aspirin ratio, we observe multimode binding, which is not limited to the outer corner region alone (Figure 3A-B). To assess structural stability, we computed the root-mean-square-deviation (RMSD) and radius of gyration (R_g_) of TD nanocage with different Aspirin ratios (Figure 3C-D). Negligible RMSD of the TD structures compared to bare TD in bulk water, indicates that Aspirin binding does not perturb the stable structural conformations of the TD, which is essential for the delivery of the Aspirin in the brain cells. We observe slightly increased RMSD and R_g_ values for the TD-100 Aspirin system, suggesting intercalation of Aspirin between two DNA base-pairs and enhanced major/minor groove bindings (Figure 3C). With the increasing TD:Aspirin ratio (i.e., 1:150) some of these interaction modes disappear due to the stacking as well as aggregation of Aspirin molecules. The size of the Aspirin dimer or multimers is larger and there is not enough space available for the Aspirin multimers to bind inside the narrow TD grooves and intercalation space. As a result, intercalation and groove bindings are less observed for the 150 Aspirin ratio, only outer corner mode bindings are dominant. This is in contrast to the more prominent groove/backbone bindings than stacked corner mode binding observed for charged aromatic molecules like dopamine, as we have seen in our previous work^19^.

**Figure 2:**
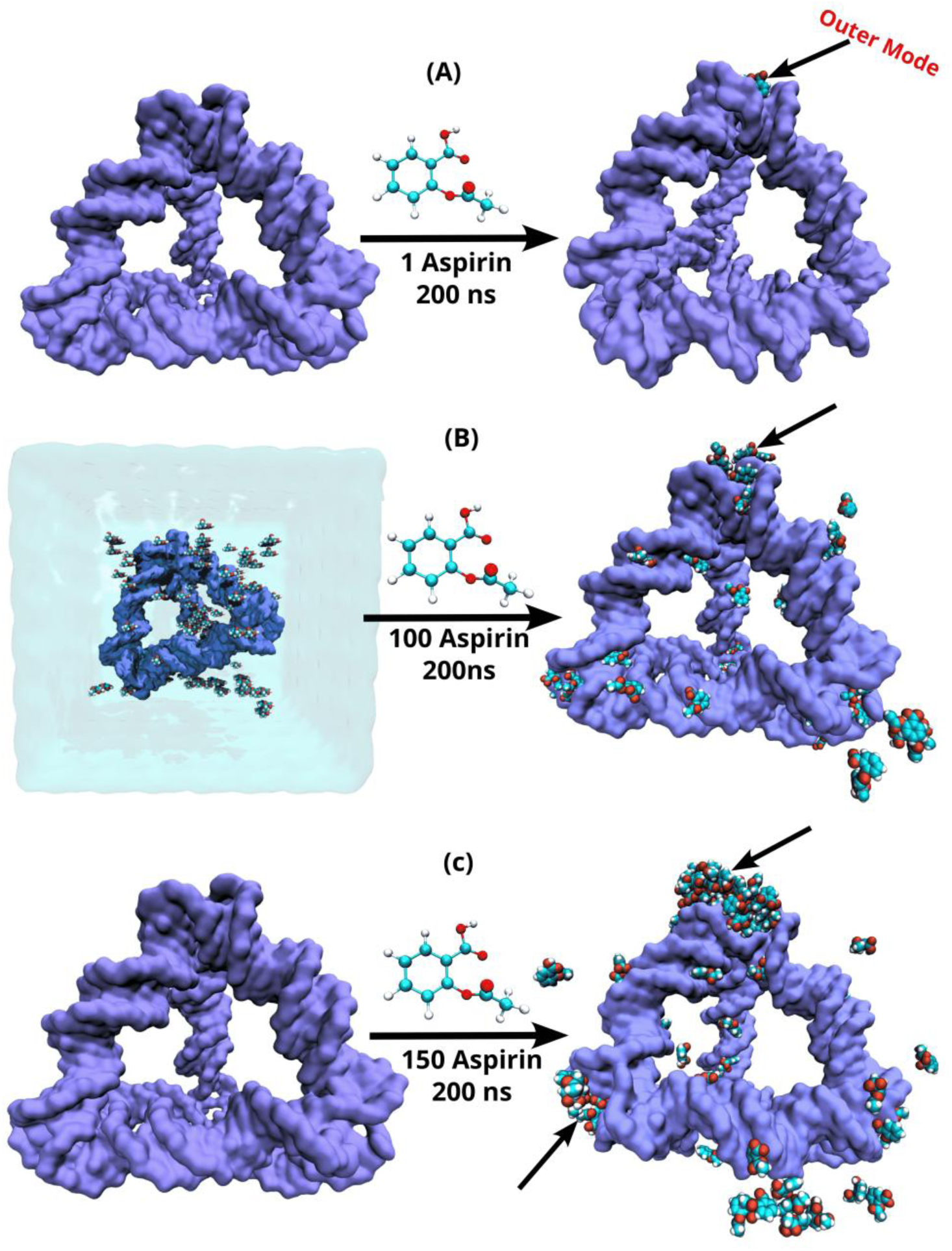
Instantaneous snapshot of Initial and final conformations of TD: Aspirin complexes at the ratios of (A)1:1, (B) 1:100, (C)1:150, from the 200 ns long atomistic MD simulations. For all three cases, outer binding modes are more prominent. In Figure (B), the simulation water box is shown in ice-blue colour and ions are not shown for clarity. The left panel are initial conformations where Aspirin molecules are placed randomly.

**Figure 3:**
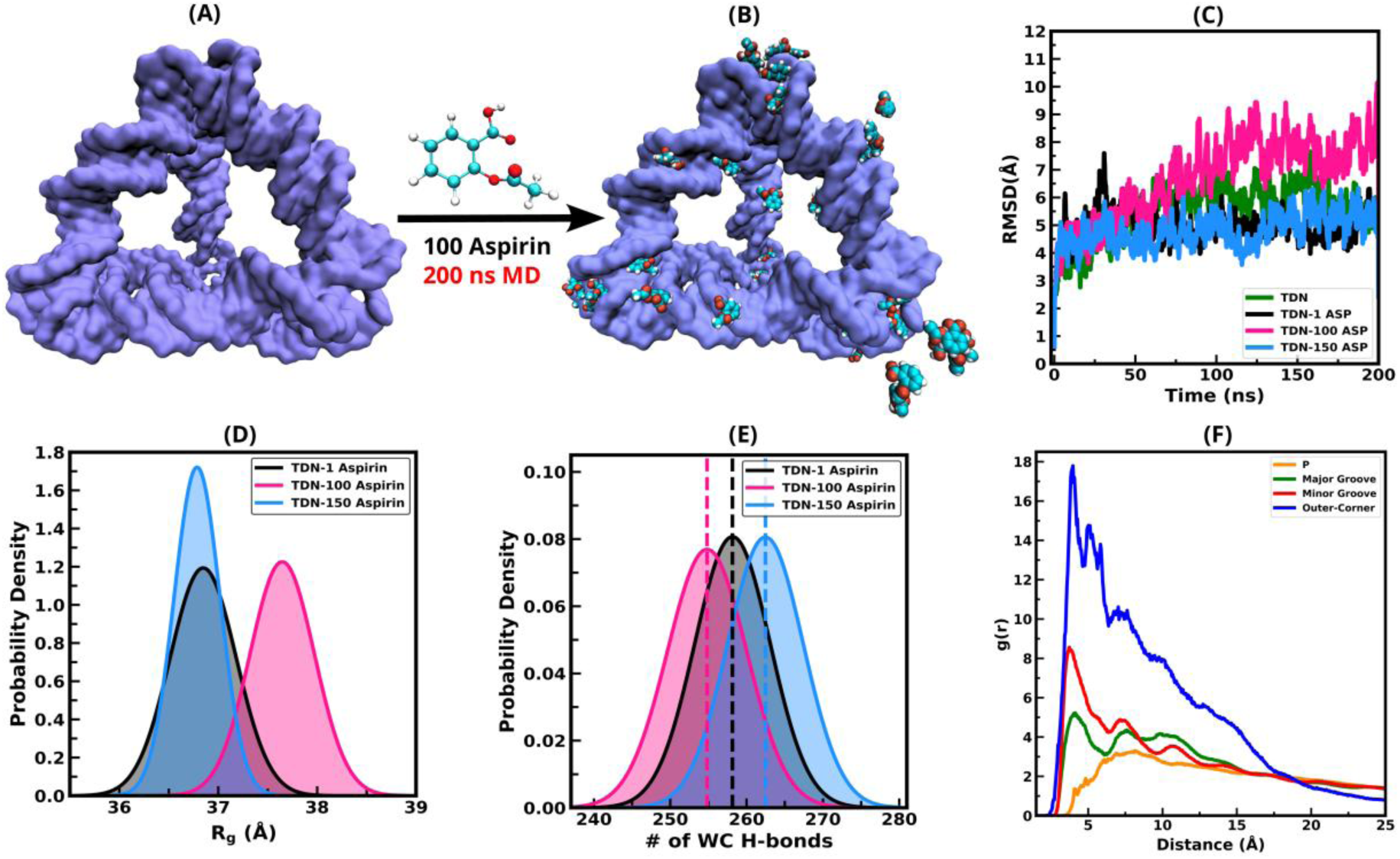
(A) instantaneous snapshot of the initial conformation of TD:Aspirin complex with 100 Aspirin molecules placed randomly in the simulation box, (B) final conformations of TD:100 Aspirin complex at the end of the 200 ns long atomistic MD simulation. binding of TD:100 Aspirin, where Aspirin shows different kinds of binding modes with TD grooves and corners. (C) root-mean-square-deviation (RMSD) for the structural stability information of TD nanocage, both in presence and absence of Aspirin molecules., (D) radius of gyration of only TD nanocage atoms in presence of 1, 100 and 150 Aspirin. (E) radial distribution function (RDF) between COM atom of Aspirin aromatic ring and the atoms from TD backbone (P) in orange, major groove atoms in red and minor groove atoms in blue. (F) number of Watson-Crick hydrogen bonds between TD base-pairs.

To investigate the structural stability of TD nanocage, we have also calculated Watson-Crick (WC) hydrogen bonds between TD base-pairs. Figure 3E shows that for a single Aspirin molecule, number of WC hydrogen bonds (258.15±3.94) remains consistent with bare TD (259.8 ±5.2), but slightly higher for 1:150 ratio (262.47±4.95), suggesting more stabilization of DNA duplex, as the stable base-pairing increases WC hydrogen bond number and for 100 Aspirin WC hydrogen bonds (254±2.18) decreases due to the disruption of base-pairing due to intercalation of Aspirin between two base pairs.

Binding modes are changing with Aspirin concentration, the radial distribution function (RDF) shows that outer corner mode is the most preferred binding sites, followed by minor, major grooves and backbones for 1:100 Aspirin ratio (Figure 3F). As a result, for the 1:100 Aspirin ratio, we observe multimodal binding, including intercalation, groove binding, backbone contacts, and both inner and outer corner stacking (Figure 4A-F). However, for the 1:150 ratio, Aspirin molecules form dimers and multimers through π-π stacking, shifting binding preference towards the outer corner mode stacking with unpaired bases, while intercalation and major/minor groove binding becomes rare.

**Figure 4:**
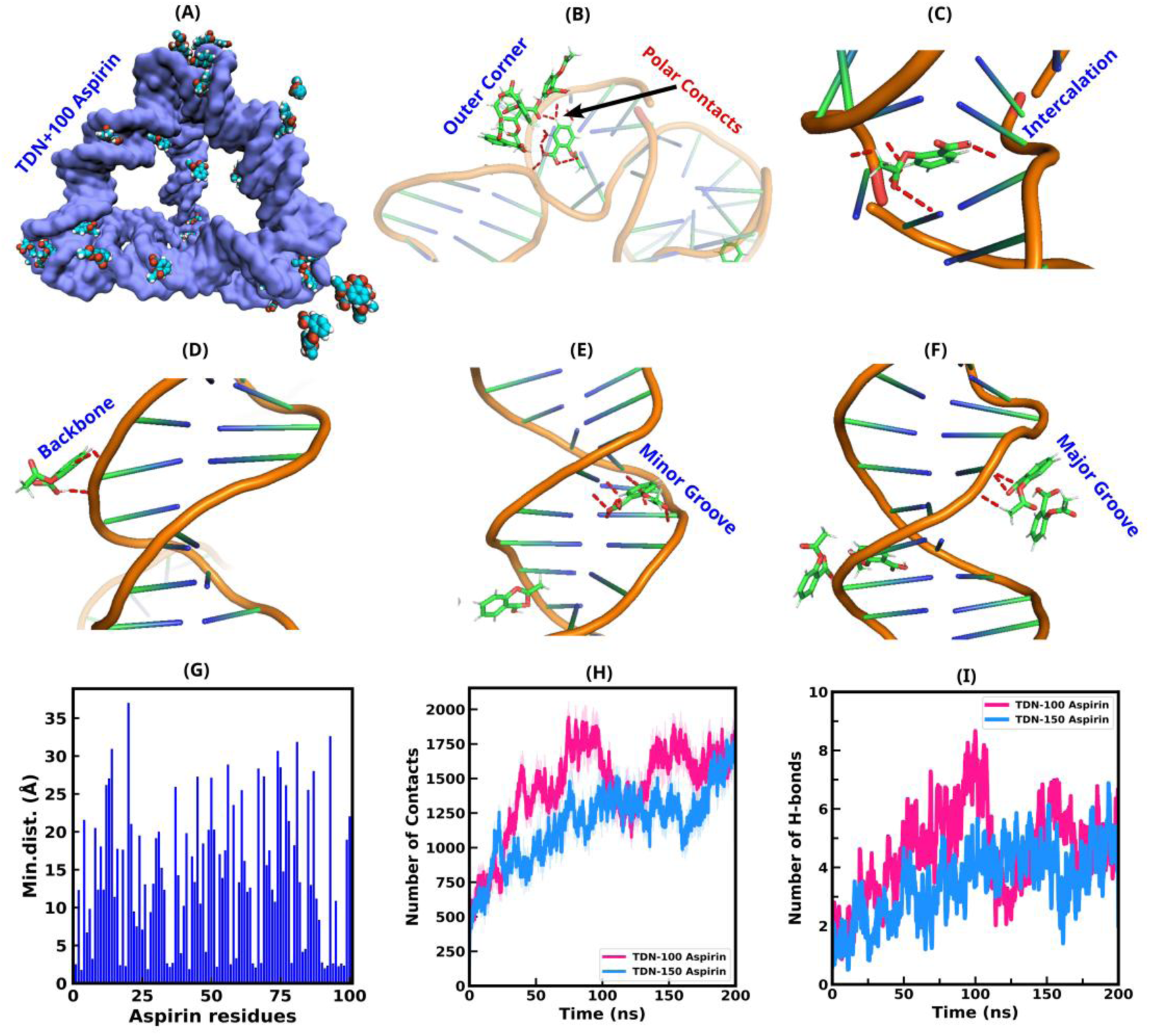
Representations of the-(A) binding modes of Aspirin in TD:100 Aspirin complex from final snapshots of 200 ns long MD simulations, (B) outer corner binding mode, (C) intercalation, (D)backbone, (E) minor Groove (N3,O2), (F) major groove (N6,O4, N4) binding modes, (G) minimum distances between each of TD nucleotides and Aspirin molecules for the 100 ratio case, (H) number of contacts formed between TD nucleotides and 100 Aspirin molecules, (I) number of hydrogen bonds formed between TD and 100 Aspirin molecules.

The minimum distance histogram shows that for the 1:100 ratio, 28 out of 100 Aspirin molecule binds (< 5Å distance) to TD nucleotides (Figure 4G) as observed from the last 20 ns of the 200 ns MD simulations. The number of contacts and hydrogen bonds formed between TD and 100 Aspirin molecules suggests that initially the number of contacts and hydrogen bonds increases and remains nearly stable for the last 50 ns, indicating that Aspirin molecules indeed show stable association with TD cages (Figure 4H-I). For the 1:150 ratio, our result shows a smaller number of contacts and hydrogen bonds formation, since there is less groove/backbone bindings and more outer corner mode stackings (Figure 4H-I).

Overall, our simulation results reveal a distinct, concentration-dependent binding mechanism for Aspirin that aligns well with our experimental observations of TD:Aspirin complexation. These findings provide a molecular-level understanding of Aspirin binding with TD nanocages supporting their potential for high-capacity loading and controlled release in brain-targeted delivery applications.

### 3.3. Aspirin entrapment efficiency in TDs and kinetic release

The percentage of the formulation’s total Aspirin that is effectively contained within the TDs is known as the entrapment efficiency. The TDs successfully contained the whole amount of Aspirin utilised, as evidenced by the achieved entrapment efficiency of 83.47%, 93.55% and 78.80% at pH 5.5, 7.4 and 9.4, respectively (Figure 5). In drug delivery systems, high entrapment efficiency is preferred because it guarantees that a larger percentage of the drug is shielded and delivered to the intended location, improving therapeutic efficacy. As shown in Figure 5(C), Release kinetics, the observation that Aspirin had a more efficient and sustained release at pH 7.4 compared to the pH 5.5 and 9.4, suggests that the TDs are designed for sustained-release targeted drug delivery. Under acidic conditions, such as those found in intracellular compartments, the TDs may undergo drug release more readily, resulting in improved drug delivery to the target site. While lowering the possibility of adverse effects on healthy tissues, this selective release improves therapeutic efficacy. Aspirin’s stability, controlled release, and improved drug release kinetics can all be improved via TDs encapsulation.

**Figure 5:**
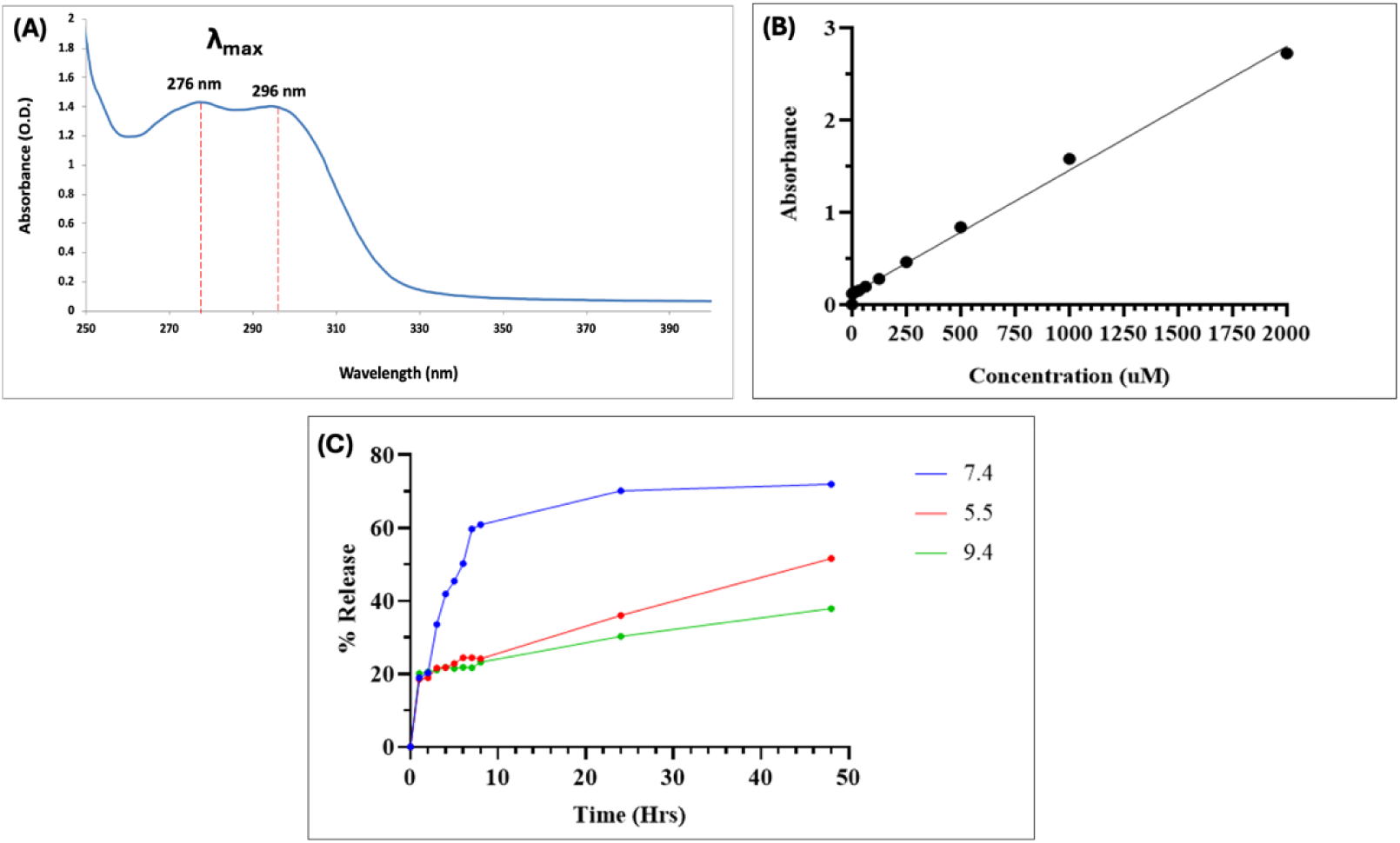
In vitro release kinetics of Aspirin from DNA Tetrahedrons at pH 5.5, 7.4 and 9.4. (A) Absorbance spectra of Aspirin (B) Standard curve of Aspirin (C) The graph shows the cumulative percentage of Aspirin released over time at pH 5.5, 7.4 and 9.4. Data represent means ± SEM of three independent experiments.

### 3.4. Morphological observation and heartbeat

The physiological effects of aspirin, TDs, and their conjugates TD were evaluated through morphological study of zebrafish larvae (3dpf): The activity, erratic swimming behaviour, mortality, latency, and deformities of aspirin (1:50, 1:100) under both normal and ischaemic conditions were carefully examined. Due to ischaemic stroke induction, all zebrafish larvae initially showed decreased locomotor activity after post-recovery treatments. This was indicative of the decreased vitality and impaired neuromuscular coordination that are frequently linked to ischaemic damage. With the exception of the untreated illness control, all treated groups showed a declining trend in latency and a progressive improvement in movement from 1 hour to 2.5 hours over time. At 1.5 and 2.5 hours, there was a discernible increase in activity, suggesting a gradual improvement in motor function. Larvae treated with TD:Aspirin (1:50) and TD:Aspirin (1:100) showed noticeably higher activities of 46.6% and 43.3%, respectively, compared to the disease control group’s low activity of 30%. The free aspirin group had a moderate initial recovery of 34.8%, but by 2.5 hours, it had drastically decreased to 20.3%, indicating rapid drug depletion and low therapeutic durability. The sustained activity levels shown in TD: Aspirin-treated groups, on the other hand, demonstrate the improved neuroprotective efficacy and regulated drug release provided by the DNA nanocage carrier technology. The TD:Aspirin (1:100) group maintained the highest locomotor activity (41.5%) by the end of 2.5 hours, indicating a dose-dependent treatment advantage, while activity in the disease control, TD-only, and TD:Aspirin (1:50) groups stayed around 32% (Figure 6b). These results suggest that aspirin conjugation with TDs enhances its bioavailability and promotes functional recovery in ischaemic zebrafish larvae by providing long-term neuroprotection.

**Figure 6:**
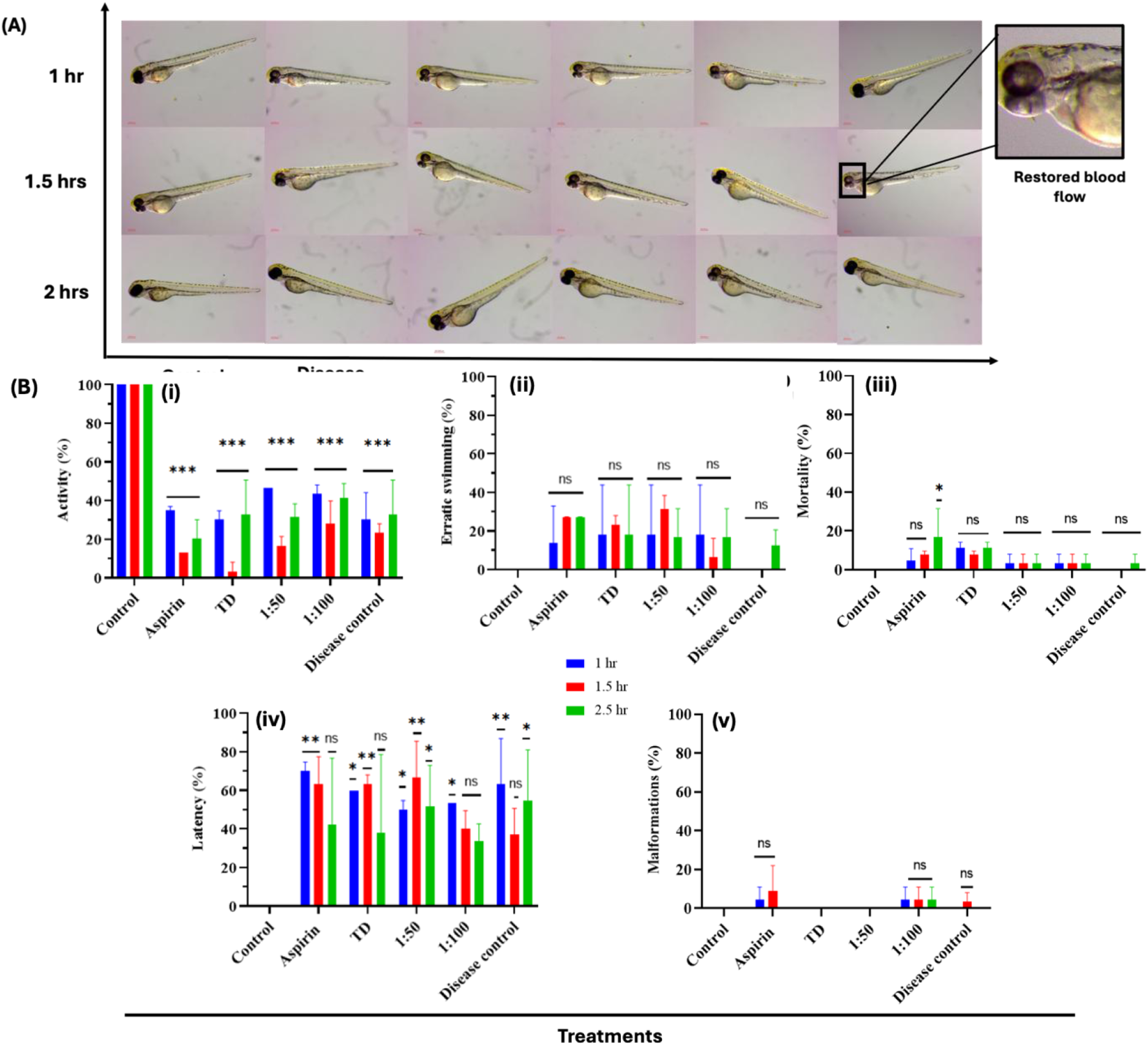
Photographs and graphs representing the morphological changes such as mortality, latency, erratic swimming, activity and malformations after inducing the ischemic stroke and post-ischemic stroke treatments in 3dpf larvae with TD, Aspirin, TD:Asp (1:50) and TD:Asp (1:100). (A) Photographs representing the various time points post-stroke treatments; 1 hr, 1.5hrs and 2 hrs with TD, Aspirin, TD:Asp (1:50) and TD:Asp (1:100). (B), (i) activity (ii) erratic swimming (iii) mortality (iv) latency (v) malformations. Data represent means ± SEM of three independent experiments. ***Statistically significant p-value (p <0.001), **Statistically significant p-value (p <0.01), *Statistically significant p-value (p < 0.05) when compared with control.

The general activity patterns seen in all treatment groups showed a strong correlation with erratic swimming behaviour. After 2.5 hours, the incidence of irregular swimming was significantly higher in larvae treated with free aspirin and TDs alone (26.9% and 18.1%, respectively) than in the disease control group (12.4%), suggesting possible neuroirritability or incomplete recovery. At 2.5 hours, however, the TD: Aspirin (1:100) group displayed a modest degree of irregular movement (16.9%), indicating better behavioural stability. Notably, irregular swimming significantly decreased in the TD:Aspirin (1:50) group, falling from 31.5% at 1.5 hours to 16.9% at 2.5 hours, indicating progressive neuromotor recovery (Figure 6b). These findings corroborate the neuroprotective role of DNA nanocages in post-ischemic recovery by indicating that their encapsulation of aspirin efficiently controls drug release and minimises behavioural impairments.

Time-dependent differences between the treatment groups were found by mortality analysis. A concentration- and time-dependent cytotoxic effect was evident in the free aspirin group, where fatality rates rose gradually from 4.5% at 1 hour to 7.8% at 1.5 hours and 16.9% by 2.5 hours. The TD-only group, on the other hand, showed comparatively steady mortality values of 11.2%, 7.8%, and 11.2% at 1, 1.5, and 2.5 hours, respectively, indicating that the nanostructures were somewhat biocompatible. Surprisingly, larvae treated with TD: Aspirin (1:50) and TD: Aspirin (1:100) showed no further increase in mortality after the first hour, maintaining a steady death rate of 3.3% throughout all time intervals (Figure 6b). This stability emphasises how DNA nanocage encapsulation protects against aspirin-induced toxicity and improves overall larval survival during ischaemic recovery.

Additionally, the TD: Aspirin (1:50) group’s latency time showed 50% at one hour and 51.5% at two and a half hours, indicating steady recovery and maintained motor response. Latency dramatically dropped in the TD: Aspirin (1:100) group from 53.33% after one hour to 33.6% at two and a half hours, indicating a steady improvement in locomotor function. In contrast to the conjugated formulations, the free aspirin group showed a decrease in latency from 70% to 42.4% within the same time period, indicating a partial but less consistent recovery. The disease control group, on the other hand, displayed a decrease in latency from 63.33% at 1 hour to 36.9% at 1.5 hours, followed by an unanticipated rise to 54.8% at 2.5 hours (Figure 6b), suggesting erratic and variable behavioural recovery. The improved and prolonged neurobehavioral recovery in TD: Aspirin-treated larvae is further supported by these latency trends, which are consistent with the comparable activity data.

The TD-only and TD: Aspirin (1:50) groups showed no morphological abnormalities, suggesting that the nanostructures were biocompatible. On the other hand, the incidence of malformations increased from 4.5% to 9.0% in the free aspirin group, indicating dose-related developmental stress. Further supporting the lower toxicity of the nanoconjugate formulation, the TD: Aspirin (1:100) group showed a low deformity rate of 4.5%, which was similar to the 3.3% seen in the disease control group.

### 3.5. Reduced cerebral thrombosis in the ischemic stroke zebrafish larva

Thrombus development in zebrafish larvae was visualised and assessed using O-dianisidine staining as a marker of ischaemic stroke induction and recovery. Blood flow restriction and clot deposition can be clearly seen because to the staining’s unique marking of hemoglobin-rich areas. Strong and distinct thrombus formation was seen close to the yolk sac area in stroke-induced larvae, indicating that ischaemia was successfully induced. On the other hand, larvae that received recovery therapies showed clearly diminished thrombus size and staining intensity, indicating better circulation and partial clot removal (Figure 7A).

**Figure 7:**
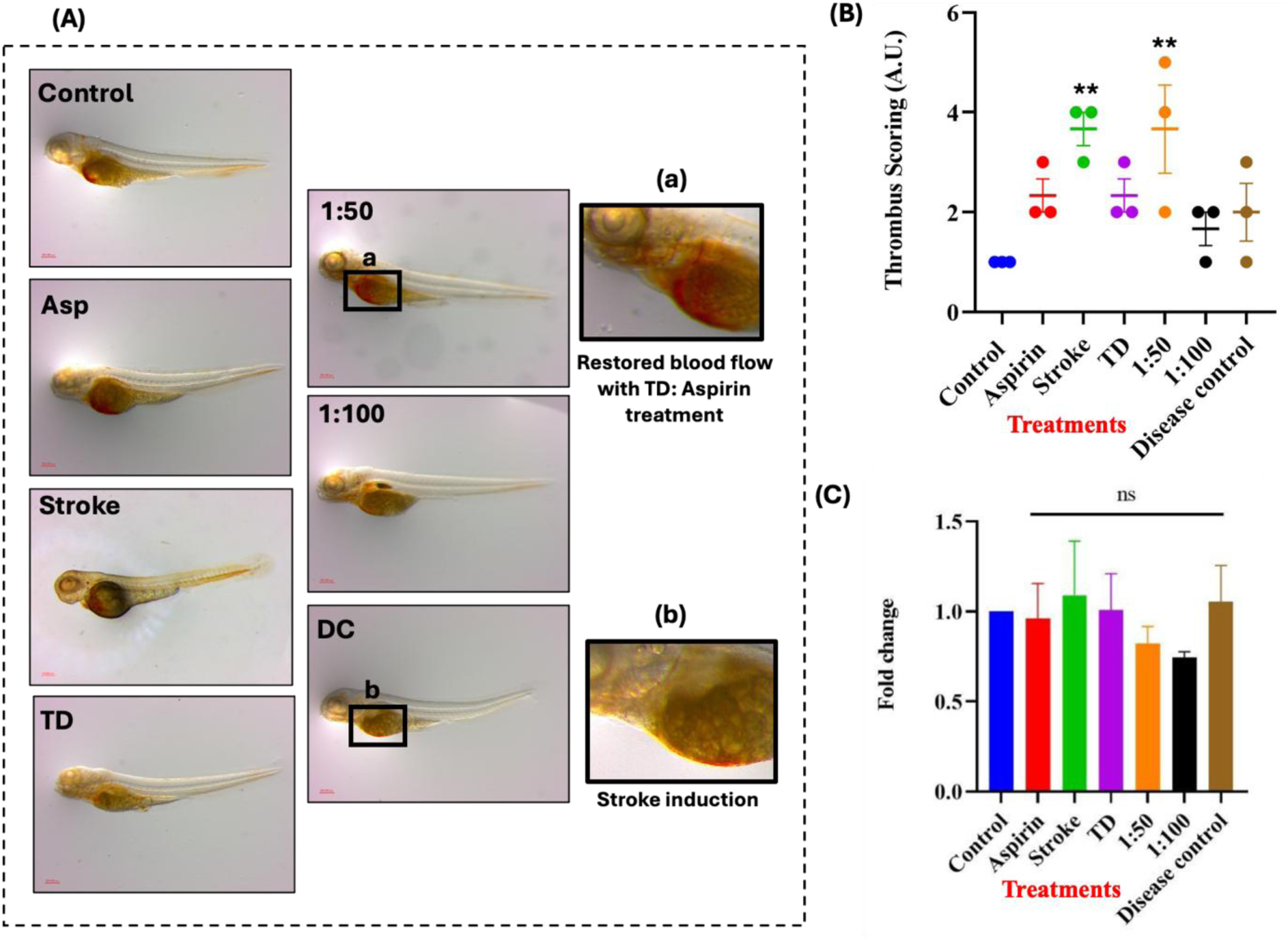
Reduced cerebral thrombosis in the ischemic stroke zebrafish larva (3dpf) treated with TD, ischemic stroke therapeutic drug Aspirin, TD:Asp (1:50) and TD:Asp (1:100). (A) Photographs representing reduced thrombosis. (B) Quantifying reduced thrombosis from image panel. (C) Heartbeats quantification of 3-dpf larvae during ischemic stroke and post-ischemic stroke treatments with TD, Aspirin, TD:Asp (1:50) and TD:Asp (1:100). Data represent means ± SEM of three independent experiments. **Statistically significant p-value (p <0.01) when compared with control.

On a scale of 1 to 5, thrombus intensity was quantitatively scored; a higher score denoted more thrombus buildup. A baseline score of 1, which indicates normal blood flow without clot formation, was obtained by the healthy control group. The ischaemic stroke group, on the other hand, had high thrombus ratings of 3–4, which indicate severe clot development and reduced perfusion. The TD: Aspirin (1:50) group demonstrated moderate thrombus scores, ranging from 3 to 5, among the recovery treatment groups, indicating a partial reduction in clot burden. Surprisingly, the TD: Aspirin (1:100) group showed considerably lower scores of 1-2, similar to the control, suggesting almost total thrombus resolution and blood flow restoration (Figure 7B).

Overall, compared to the stroke group, the larvae receiving recovery therapy showed a significant decrease in thrombus size and intensity, as shown by the O-dianisidine staining results. The significant decrease in thrombus formation in the TD:Aspirin (1:100) group indicates that Aspirin’s antithrombotic activity is enhanced by the higher ratio of DNA nanocage conjugation, most likely through better bioavailability and regulated release. These data demonstrate that TD:Aspirin nanoconjugates successfully assist thrombus breakdown and improve vascular healing following ischaemic injury in zebrafish larvae.

The ischaemic stroke-treated larvae showed a 1.08-fold shift in heart rate, which is suggestive of increased physiological stress after ischaemic induction. By contrast, the heart rate fold changes in the TD-only (1.00) and free aspirin (0.96) groups were comparable to the disease control (1.00), indicating that these therapies did not worsen cardiac stress. When taken as a whole, our results demonstrate the safety and biocompatibility of TD:Aspirin formulations in preserving normal heart growth and function throughout ischaemic recovery. Additionally, compared to the stroke-induced group, the recovery treatment groups’ heart rates were found to be slower, suggesting less physiological stress. In particular, the heart rates of the TD:Aspirin (1:50) and TD:Aspirin (1:100) groups changed by 0.82 and 0.74 times, respectively, in comparison to the disease control, indicating better cardiac stability and reduced stress levels during recovery (Figure 7C).

### 3.6. Intracellular reactive oxygen species

DCFDA staining was used to measure intracellular ROS levels, and ImageJ software was used to quantify the fluorescence intensity. ROS levels were significantly elevated in stroke-induced larvae, almost eight times greater than in control larvae, indicating oxidative stress linked to ischaemic injury. Compared to the ischaemic group, larvae treated with TD: Aspirin (1:50) showed a fourfold rise, whilst other recovery therapy groups showed roughly a threefold increase, suggesting a considerable decrease in ROS formation (Figure 8A,B). The reduction of oxidative stress in TD: Aspirin-treated larvae indicates that the DNA nanocages improve the antioxidant and neuroprotective properties of aspirin in addition to facilitating its controlled and prolonged release. These results suggest that the nanoconjugate formulation is more effective than free medication delivery at reducing ischemia-induced cellular damage. Increased cellular healing and functional restoration would probably result from prolonging the course of treatment, which would further normalise ROS levels towards control values.

**Figure 8:**
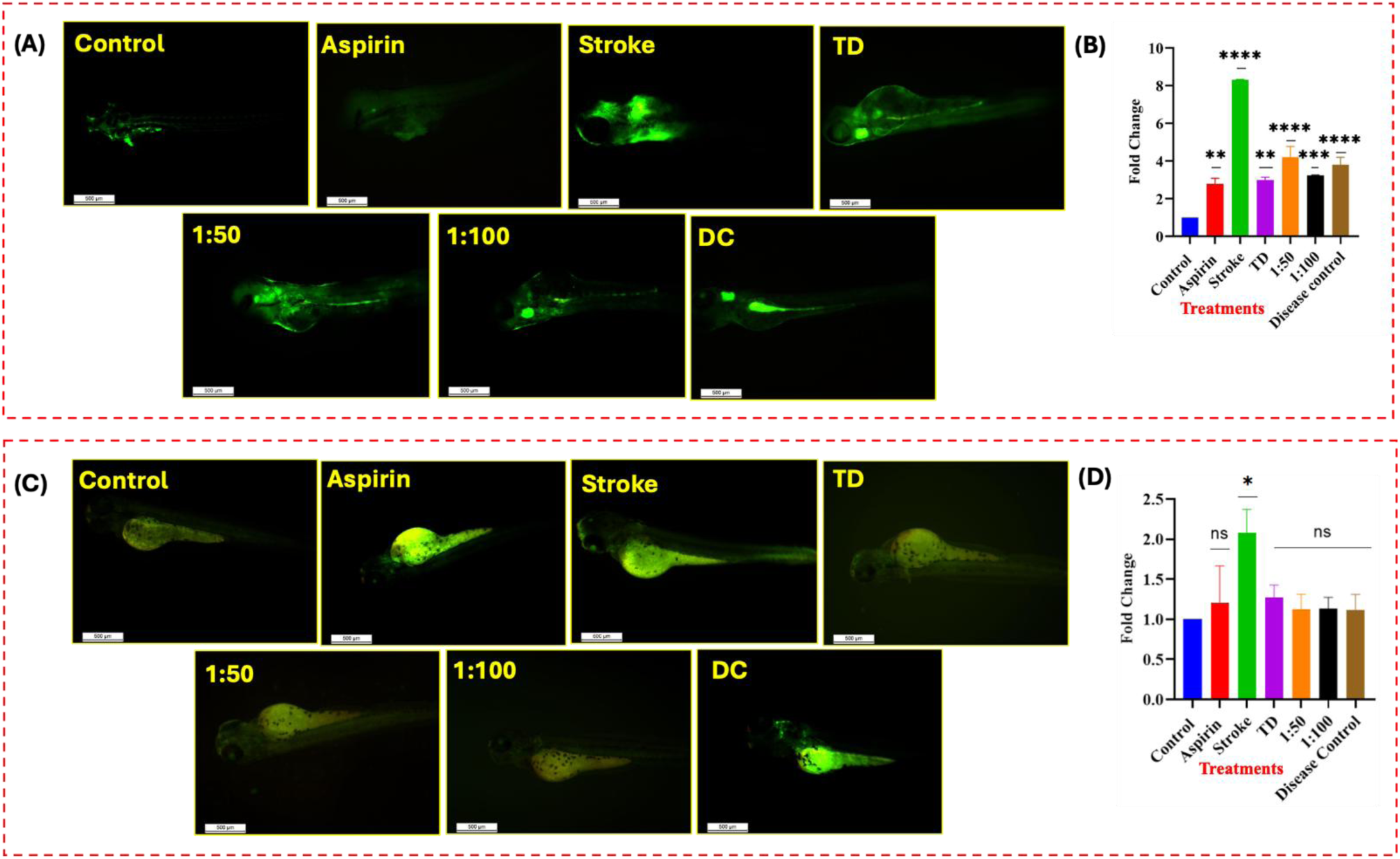
(A) Intracellular ROS generation detected by DCFDA dye. 3dpf larvae were exposed to TD, Aspirin, TD:Asp (1:50) and TD:Asp (1:100) post-ischemic stroke. (B) Bar graph depicts fold change in ROS generation. (C) Acridine orange staining of 3dpf post-ischemic stroke larvae for apoptotic exhibition. (D) Bar graph depicts fold change in apoptosis events. Data represent means ± SEM of three independent experiments. Compared with the control group: *P < 0.05, **P < 0.01, and ***P < 0.001. ****P < 0.0001 when compared with control.

### 3.7. Apoptosis

Apoptosis, also known as programmed cell death, is a characteristic of ischaemic injury and is strongly linked to increased intracellular ROS levels. Apoptotic cell counts in stroke-induced larvae almost doubled in comparison to controls, indicating the ischemia-induced oxidative stress and neuronal injury. On the other hand, larvae treated with TD:Aspirin and other recovery formulations showed a slight ∼1.3-fold increase in apoptosis, which was similar to control levels but not statistically significant according to ANOVA analysis. This decrease in apoptosis is consistent with the observed diminution of ROS formation, indicating that aspirin delivered via DNA nanocage efficiently reduces oxidative damage. The neural crest region was the primary location of apoptotic cells, which were visible as green fluorescent spots under blue light, underscoring the region’s susceptibility to ischaemic stress. A brief ∼1.5-fold increase in apoptosis was first seen in all recovery groups, most likely as a result of lingering ischaemic effects. However, apoptosis levels dramatically dropped after 2.5 hours of therapy, suggesting active cellular regeneration and recovery (Figure 8C,D). These results highlight TD:Aspirin’s dual effect in lowering oxidative stress and safeguarding brain cells, which in turn promotes neuroprotection and functional recovery in zebrafish larvae that have had strokes.

### 3.8. Reduced brain calcification

Calcium accumulation in the form of punctate nodules is a key indicator of NMDAR dysfunction and excitotoxicity, commonly observed following ischemic stroke^54^. No calcium puncta were found in the control larvae, suggesting balanced intracellular calcium levels and appropriate NMDAR function. In contrast, stroke-induced larvae exhibited significant calcium deposition, with 74 vivid green puncta visualised, demonstrating substantial NMDAR dysfunction and intracellular calcium overload. Recovery treatment considerably decreased this effect: both the Aspirin and TD-alone groups showed 7 and 6 puncta, respectively, whereas the TD: Aspirin (1:50) group exhibited just 3 puncta, showing its greater efficacy in restoring calcium homeostasis. There were two puncta in the disease control group and six in the TD: Aspirin (1:100) group. These results demonstrate that TD: Aspirin formulations, particularly at a 1:50 ratio, significantly reduce excitotoxic calcium buildup, possibly through sustained drug release and increased neuroprotection (Figure 9). By limiting calcium overload, the treatment preserves neuronal function and contributes to improved recovery following ischemic injury in zebrafish larvae.

**Figure 9:**
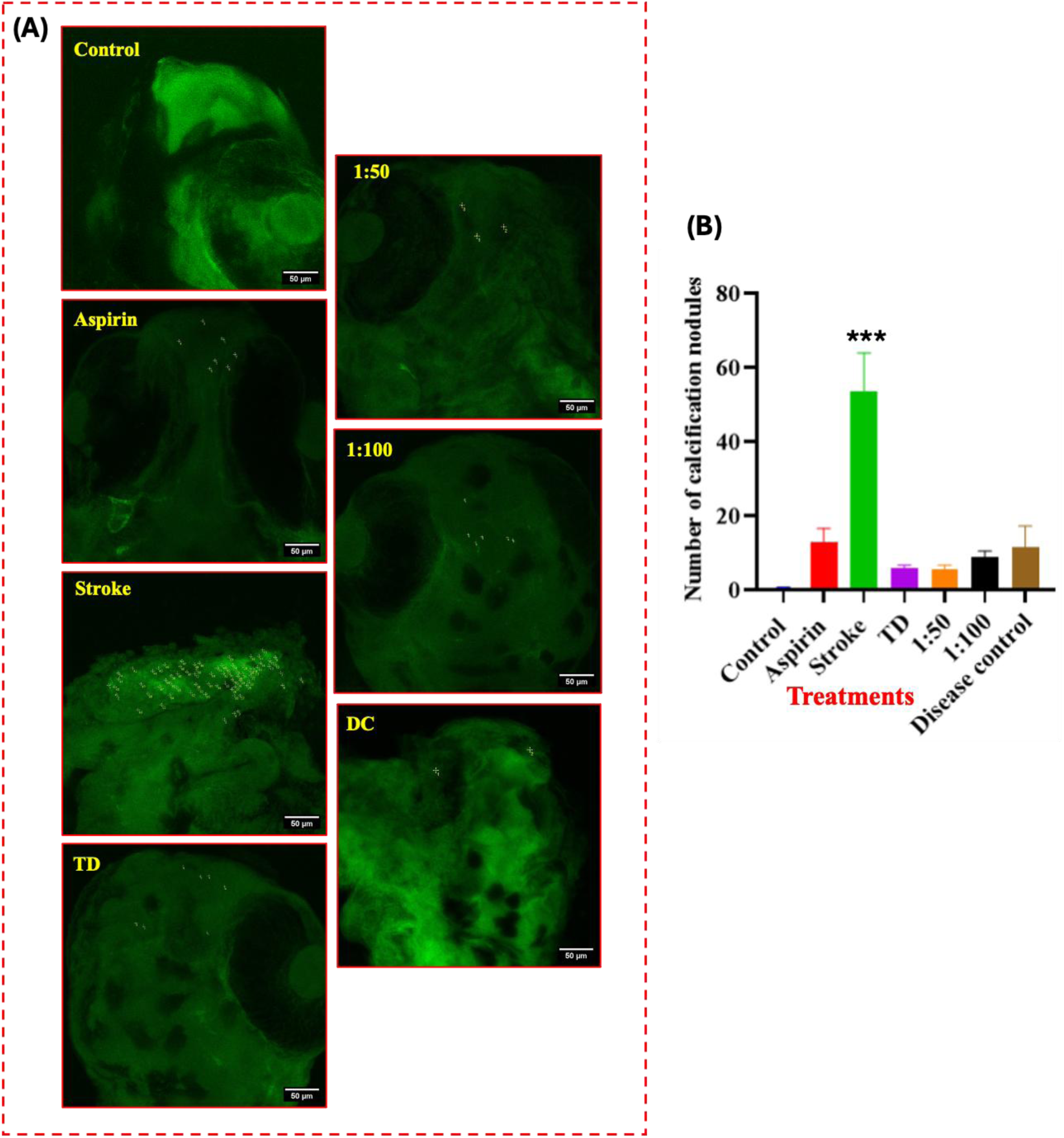
(A) Brain calcification stained by calcein dye. 3dpf larvae were exposed to TD, Aspirin, TD:Asp (1:50) and TD:Asp (1:100) post-ischemic stroke. (B) Bar graph depicts fold change in calcification nodules. ***P < 0.001. when compared with the control.

### 3.9. Gene Expression

#### 3.9.1. Specific genes associated with ischemia

An essential growth factor, brain-derived neurotrophic factor (BDNF) improves neuronal function in stressful situations. N-methyl-d-aspartate receptor (NMDAR)-triggered excitotoxicity, dendritic regeneration, and anti-apoptotic effects via B-cell lymphoma 2 (Bcl-2) protein or apoptosis-related proteins like Bad and Bim are some of the neuro-protective effects of BDNF.^55^. BDNF expression was significantly elevated in stroke-induced larvae, exhibiting a 95.0-fold increase in comparison to control, most likely as a compensatory reaction to ischemia-induced neuronal stress. In contrast, recovery treatments considerably modified BDNF levels, indicating reduction of cellular stress and enhancement of functional recovery. The TD: Aspirin (1:100) group exhibited a 17.2-fold increase, while TD-only and TD: Aspirin (1:50) treatments showed 37.4- and 27.4-fold increases, respectively. Notably, the free Aspirin group displayed a comparatively modest 5.1-fold increase (Figure 10A). These findings suggest that DNA nanocage-mediated delivery of Aspirin fine-tunes BDNF expression, reducing excessive stress responses while supporting neuroprotective pathways. This controlled modulation likely contributes to enhanced neuronal survival, improved synaptic function, and accelerated recovery in zebrafish larvae following ischemic insult. In case of an impaired signalling of BDNF, it can lead to various neurologic, psychiatric disorders like - stroke, epilepsy, and trauma. Along with this some of the neurodegenerative diseases like Alzheimer’s disease (AD), Parkinson’s disease (PD), and Huntington’s disease (HD) ^55^.

**Figure 10:**
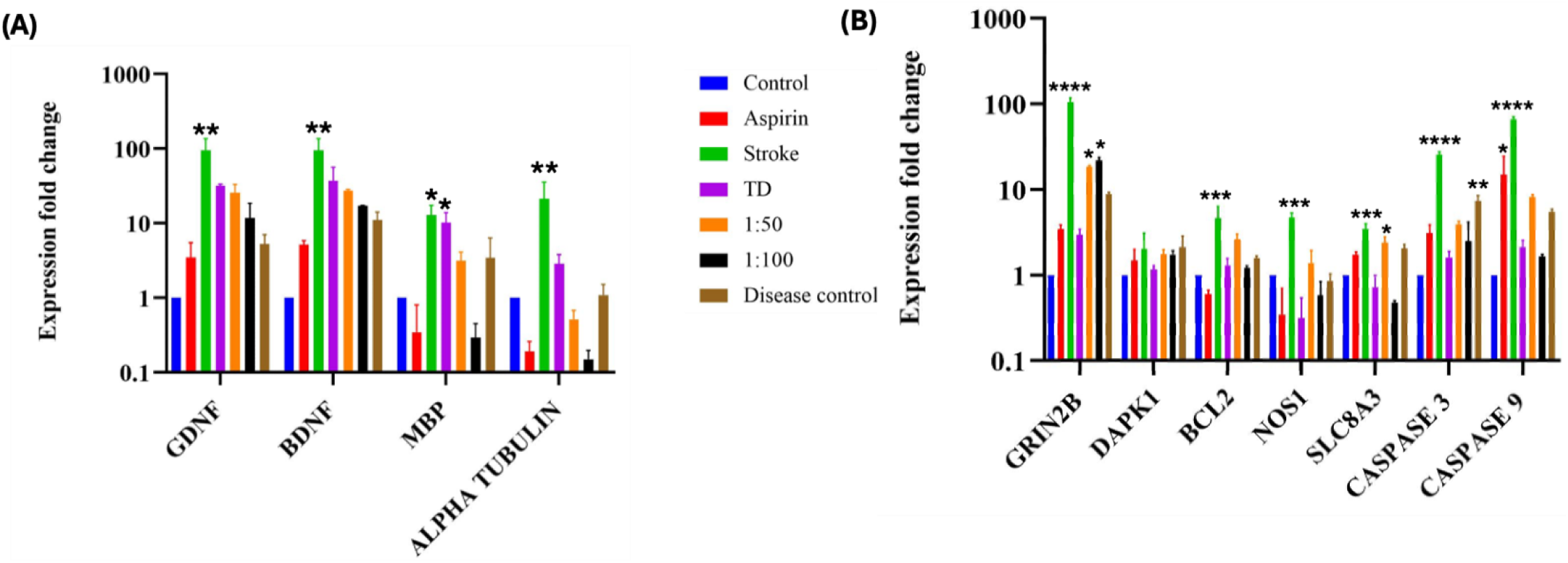
Bar graph depicting fold change in relative gene expression of zebrafish embryos exposed to TD, Aspirin, TD:Asp (1:50) and TD:Asp (1:100) post-ischemic stroke. Data represent mean±SEM of three independent experiments. *P < 0.05, **P < 0.01, and ***P < 0.001. ****P < 0.0001 when compared with control. (A) Important genes involved in the ischemic stroke cascade. (B) Excitotoxicity pathway genes.

Glial cell-derived neurotrophic factor (GDNF) was first isolated from the glioma cell line of rats. It was found that for dopaminergic neurons, GDNF was a potent survival neurotrophic factor. In later studies, the protective effect of GDNF was also seen in other neuronal populations, like non-adrenergic and motor neurons. In CNS, microglia and astrocytes mainly release GDNF, once released it binds to GFRα1 in the neuronal membrane. The binding can trigger a downstream signalling pathway. The GDNF-GFRα1-RET signalling pathway has been shown to take part in neuronal development, differentiation, and survival under both healthy and diseased conditions^56^. Glial cell line-derived neurotrophic factor (GDNF) expression was markedly upregulated in the stroke group, exhibiting a 94.1-fold increase relative to control, reflecting a strong compensatory response to ischemic injury. Treatment with TD: Aspirin (1:100) significantly attenuated this upregulation, reducing GDNF expression to an 11.7-fold increase. In comparison, the TD-only and TD: Aspirin (1:50) groups showed moderate increases of 32- and 25.5-fold, respectively. The free Aspirin group exhibited a mild 3.5-fold increase, while the disease control group showed a 5.2-fold change relative to control. These results indicate that DNA nanocage-mediated delivery of Aspirin effectively modulates GDNF expression, preventing excessive stress-induced upregulation while supporting neuroprotective and regenerative pathways in zebrafish larvae following ischemic insult. Deletion of GDNF in astrocytes increased oxidative stress, and a decreased proliferation of reactive astrocytes was seen in the peri-infarct region, along with reduced neurogenesis in the dentate gyrus^56^.

Myelin Basic Protein (MBP) is responsible for attaching the myelin sheath to oligodendrocytes. In a study, it is shown that serum levels of MBP increase in the early period after acute ischemic stroke^57^. It is an important structural component in the central and peripheral nervous system for myelin sheath formation. It is considered to be a marker of demyelination in brain injury and during chemical exposure^58^. Myelin basic protein (MBP) is a critical marker of myelin integrity and neuronal function. In stroke-induced larvae, MBP expression was markedly upregulated, showing a 12.8-fold increase relative to control, likely reflecting stress-induced dysregulation of myelination following ischemic injury. Additionally, the TD-only group showed increased expression (10.3-fold), indicating that myelin disruption cannot be completely prevented by the nanostructures alone. The TD: Aspirin (1:50) and disease control groups, on the other hand, showed moderate increases of 3.1 and 3.4 times, respectively, indicating partial normalisation. Surprisingly, MBP expression was significantly reduced in the TD: Aspirin (1:100) group to 0.2-fold, whereas free aspirin treatment produced a similar decrease of 0.3-fold in comparison to control. This MBP level normalisation indicates that higher-ratio TD: Aspirin therapy successfully stabilises myelin structure, lessens dysregulation brought on by stress, and encourages functional neuronal recovery. These findings highlight the potential of DNA nanocage-mediated Aspirin delivery to preserve myelin integrity and support neuroprotective outcomes in post-ischemic zebrafish larvae.

Acetylated α-tubulin is essential for maintaining the structure and function of mature dendritic spines in brain. Upregulation of the acetylated α-tubulin results in protection of dendritic spines in penumbra area after stroke. This is essential for neuronal function and connectivity in the regions surrounded by the infarct. The reduction in the acetylated α-tubulin showed an aggravated dendritic spine injury, worsened motor dysfunction post-ischemic stroke. This proves that acetylation of α-tubulin is known to have a protective shield against ischemic damage^59^. In stroke-induced larvae, α-tubulin expression increased dramatically, by 21.5-fold relative to control, reflecting cytoskeletal disruption and compensatory remodelling in response to ischemic injury. The TD-only and disease control groups exhibited modest increases of 2.8- and 1.09-fold, suggesting partial stabilization of cytoskeletal structure. Interestingly, the TD: Aspirin (1:50) group showed a decrease to 0.5-fold, while free Aspirin and TD: Aspirin (1:100) treatments exhibited pronounced reductions to 0.19- and 0.15-fold, respectively. These results indicate that TD: Aspirin, particularly at a higher ratio, effectively normalizes α-tubulin expression, stabilizes the neuronal cytoskeleton, and mitigates ischemia-induced structural stress. This cytoskeletal stabilization likely contributes to enhanced neuronal integrity and functional recovery in post-ischemic zebrafish larvae.

#### 3.9.2. Exploring the excitotoxicity pathway

An ischemic cascade is initiated that leads to intensive neuronal injury due to ischemic stroke and impaired blood flow. A failure of high-energy metabolism in neural cells due to reduced blood flow causes the blockage of ATP synthesis. This results in the impaired activity of ion pumps. The NMDA receptors get inhibited, causing an intracellular Ca^2+^, Na^+^, and Cl^-^accumulation^54^. Due to ATP reduction, there is lysis of mitochondria and lysosomes, hydrolysing intracellular material and further causing calcium influx. Excitotoxicity is a result of the release of the excitatory amino acid, glutamate due to increased calcium concentrations^60^. This also activates the enzyme nitric oxide synthase, and calcium-dependent enzymes like protein kinase C and phospholipase A2, aggravating cell death ^61^. Furthermore, there is a release of free radicals, reactive oxygen species (ROS), and danger-associated molecular patterns (DAMPs) by the injured neural cells. This leads to the activation of microglia and astrocytes, resulting in the release of inflammatory factors. The release of inflammatory factors causes the recruitment of leukocytes and white blood cells, further causing adherence to endothelial cells. This leads to the alteration of the endothelial cytoskeleton and degeneration of pericytes. Further affecting the endothelial junction proteins (adherens junctions, gap junctions, and tight junctions), which leads to the disruption of the blood-brain barrier (BBB). The compromised BBB leads to pro-inflammatory cytokines accumulation in the injured site, worsening neural cells and oligodendrocyte death^62^. The genes majorly involved in this cascade are GRIN2B (NMDA receptor function), SLC8A3 (affects the sodium calcium channels), NOS (Nitric oxide synthase regulation), DAPK1 (death-associated protein kinase), Bcl 2 (anti-apoptotic), caspase 3, and caspase 9 (pro-apoptotic).

The N-methyl D-aspartate receptor (NMDAR), an ionotropic receptor, is made up of various subunits. In a study done by Liu and his coworkers on the rat model of focal ischemic stroke, they observed that the activation of synaptic/extra-synaptic NR2A and NR2B containing NMDAR resulted in the opposite effects on neurons. The NR2A-containing2 NMDAR is known to have a protective action, promoting survival of neurons, while the NR2B-containing NMDAR is known to enhance excitotoxicity, thereby increasing apoptosis of neurons^63^. NMDAR activation leads to excessive influx of extracellular calcium and an imbalance of cellular ion homeostasis^64^. The NMDAR-NR2B is also known as GRIN2B. This subunit is found to be included in the development of synapses, macromolecular organization, actin cytoskeleton, and plasticity^65^. In stroke-induced larvae, GRIN2B expression was dramatically upregulated, showing a 105.2-fold increase relative to control, reflecting heightened excitotoxic stress and dysregulated calcium influx associated with ischemic injury. Treatment with TD: Aspirin at 1:50 and 1:100 ratios substantially moderated this overexpression, with fold changes of 18.6 and 21.8, respectively, suggesting partial restoration of NMDAR function and intracellular calcium homeostasis. While free aspirin and TD-only treatments showed smaller increases of 3.4- and 2.9-fold, respectively, the disease control group showed a moderate 8.8-fold increase (Figure 10B). These findings suggest that TD: Aspirin administration successfully reduces excitotoxicity and increases neuronal lifespan by attenuating GRIN2B overactivation. In zebrafish larvae after ischaemic injury, this modulation probably helps with neuroprotection, better calcium regulation, and improved functional recovery. Knockout of this subunit leads to lethality, and hence its precise function is comparatively difficult to access^66^.

Death-Associated Protein Kinase 1 (DAPK1) is essentially a Ca2+/calmodulin-dependent serine/threonine-protein kinase. DAPK1 phosphorylation leads to apoptotic cell death. In an ischemic event, overactivation of NMDAR leads to increased Ca2+ influx, thus activating Ca2+/calmodulin^4^. This activation leads to the stimulation of calcineurin phosphatase, resulting in dephosphorylation and activation of DAPK1^67^. The ischemic injury is further aggravated by the transfer of activated DAPK1 to the GluN2B subunit of NMDARs^68^. DAPK1 expression remained relatively consistent across all treatment groups. The highest upregulation was observed in the disease control group, showing a 2.1-fold increase relative to control, while the stroke group exhibited a similar 2.01-fold increase. TD: Aspirin treatments at both 1:50 and 1:100 ratios displayed moderate increases of 1.75-fold compared to control. Other treatment groups showed lower expression levels: the TD-only group exhibited a 1.15-fold increase, and the free Aspirin group displayed the lowest upregulation at 1.48-fold. These results indicate that DAPK1 expression is only modestly affected by stroke and treatment interventions, with TD: Aspirin formulations maintaining expression levels closer to baseline, suggesting minimal impact on DAPK1-mediated apoptotic pathways in zebrafish larvae. Disruption of the interaction between NMDAR and DAPK1 has shown reduced neuronal excitotoxicity in mouse IS models and was able to suppress the NMDAR current in vitro^68^. DAPK1 dephosphorylation is prevented by inhibiting either NMDAR or calcineurin. Inhibition of DAPK1 leads to protection against ischemic injury when tested in both cultured neurons and in vivo. Hence, the potential treatments for treating ischemic stroke could be formed on the basis of DAPK1 inhibition^69^. In another study, the GluN2B-DAPK1 interaction was interrupted via an interference peptide Tat-NR2B-CT. This peptide contained the C-terminal phosphorylation site of GluN2B. thereby, mitigating the DAPK1-induced phosphorylation of GluN2B-containing receptors, blocking calcium influx. This interruption was shown to provide neuroprotection against excitotoxicity in both - in vivo and in vitro ischemic models ^70, 71^.

Neuronal nitric oxide synthase (nNOS) is activated by excessive influx of calcium. Activated GluN2B receptors are responsible for the translocation of GluN2B conjugated nNOS from the cytoplasm to the membrane. This results in the production of excess neurotoxic nitric oxide^70^. In later studies, it was shown that a scaffolding protein called postsynaptic density-95 protein (PSD-95) is responsible for binding of nNOS to C-terminal of GluN2B, thereby bringing nNOS in close proximity to NMDARs, allowing enough calcium to activate nNOS ^72, 73^. NOS1 expression was markedly upregulated in the stroke group, showing a 4.6-fold increase relative to control, reflecting enhanced nitric oxide signalling associated with ischemic stress. In contrast, the TD: Aspirin (1:50) group exhibited a modest 1.3-fold increase, comparable to control levels, indicating effective normalization of NOS1 expression. Both the free Aspirin and TD-only groups showed significant decreases to 0.34- and 0.31-fold, respectively, while the TD: Aspirin (1:100) group demonstrated a 0.57-fold reduction. These findings suggest that TD: Aspirin treatment helps restore NOS1 expression toward physiological levels, potentially mitigating oxidative and nitrosative stress and contributing to neuroprotection in zebrafish larvae following ischemic injury. Interrupting the NMDAR/PSD-95/nNOS interaction with modified drugs such as Tat-NR2B9c, IC87201, and ZL006 was able to prevent the nNOS activation via GluN2B receptors. This intervention resulted in decreased in vitro neuronal damage, and reduced in vivo infarct volume and neurological deficits ^73, 74^.

NCX reversal is seen during neuronal ischemia because of ionic disturbances. This leads to increased calcium levels intracellularly. Upon reperfusion, the NCX may regain the forward mode^75^. High calcium influx is responsible for inducing several downstream cascades of events, responsible for cell death, like: protease, lipase, nuclease, and caspase triggering. Na+/Ca2+ exchanger, also known as NCX, is a membrane antiporter that is responsible for the efflux of calcium in exchange of sodium influx^64^. The NCX plays an important role against the increasing accumulation of sodium and calcium ions^76^. In cerebral ischemic events, it is shown that an increase in the activity of NCX is shown to promotes neuronal survival^64^. When upregulated, in cerebral ischemic models, NCX restores the sodium/calcium homeostasis, thereby reducing the neurological damage^77^. SLC8A expression exhibited a pattern similar to that of NOS1. In stroke-induced larvae, SLC8A was upregulated 3.4-fold relative to control, reflecting altered calcium handling under ischemic stress. The TD: Aspirin (1:50) and disease control groups showed comparable moderate increases of 2.3- and 2.04-fold, respectively. The free Aspirin group displayed a 1.7-fold increase, whereas the TD-only group exhibited a 0.7-fold decrease. Notably, the TD: Aspirin (1:100) group demonstrated a more pronounced reduction to 0.48-fold relative to control. These results indicate that higher-ratio TD:Aspirin treatment effectively normalizes SLC8A expression, potentially stabilizing intracellular calcium homeostasis and contributing to neuroprotection in zebrafish larvae following ischemic injury. In a study by Boscia et al., they demonstrated that on silencing the NCX3 gene, there was reduced myelination observed because of defective oligodendrocyte differentiation and maturation^78^.

#### 3.9.3. Genes associated with apoptosis

The anti-apoptotic proteins like Bcl-2 and Bcl-xL provide protection to the outer mitochondrial membrane (OMM) integrity and also prevent the formation of cytochrome C release, thereby preventing the initiation of apoptotic pathways^79^. The increased calcium levels can initiate the intrinsic apoptotic pathways through calpain activation and also can stimulate the opening of the mitochondrial permeability transition pore. Activated calpains and caspase 8 are responsible for truncating Bid. The truncated Bid can translocate to mitochondria and counteract the antiapoptotic proteins like Bcl-2, and is able to interact with proapoptotic proteins. As a result, the mPTP is opened and cytochrome c, along with other factors, is released into the cytoplasm. In the presence of dATP, the apoptosome complex is formed, which consists of cytochrome c, Apaf-1, and procaspase-9. This apoptosome can activate caspase 9 and further activate caspase-3 downstream^70^. Generally, higher expressions of anti-apoptotic genes correlate with low expression of apoptotic genes. Multiple studies have shown that ROS-mediated cell death can be analysed by the expression of apoptotic genes in zebrafish^50^. Bcl-2, an antiapoptotic gene, was upregulated in the TD: Aspirin (1:50) group, which exhibited a 2.6-fold increase. The TD-only and TD: Aspirin (1:100) groups displayed smaller fold changes of 1.3- and 1.2-fold, and free Aspirin alone showed a reduction to 0.6-fold relative to control. The Bcl-2 protein family consists of proapoptotic proteins such as Bax, Bid, and p53-upregulated modulator of apoptosis (PUMA) and antiapoptotic proteins such as Bcl-248. Caspase 3, a key executioner of apoptosis, was markedly upregulated in the stroke group, exhibiting a 25.5-fold increase relative to control, indicating extensive apoptotic activity following ischemic injury. The disease control group also showed significant upregulation at 7.3-fold. In recovery treatment groups, caspase 3 expression was more moderate: the TD-only and TD: Aspirin (1:100) groups showed 1.6- and 2.5-fold increases, respectively. The expression levels of free aspirin and TD:aspirin (1:50) treatments were 3.1 and 3.9 times greater, respectively, but still within control. These findings imply that TD: Aspirin formulations, especially at higher ratios, successfully regulate caspase 3 activation, reducing apoptosis and fostering neuroprotection in zebrafish larvae after ischaemic injury. The apoptotic pathway was extensively activated after ischaemic injury, as evidenced by the 66.7-fold rise in caspase 9, an activator of apoptosis, in stroke-induced larvae. With a 14.9-fold upregulation in comparison to control, the free aspirin group likewise showed a significant increase. A considerable 8.2-fold rise was seen in the TD: Aspirin (1:50) group, whereas a lesser 2.1-fold upregulation was seen in the TD-only group. Interestingly, the TD: Aspirin (1:100) group showed a slight 1.6-fold increase in comparison to the control, suggesting that caspase 9 activation was effectively attenuated (Figure 10B). These findings imply that higher-ratio TD: Aspirin therapy can successfully regulate apoptotic signalling, enhancing neuroprotection and neuronal survival in zebrafish larvae following ischaemic injury.

## 4. Conclusion

In a zebrafish model of ischaemic stroke, we examined the therapeutic potential of DNA tetrahedron (TD)-mediated aspirin administration by evaluating behavioural, morphological, and molecular results. Reduced locomotor activity, increased latency, elevated reactive oxygen species (ROS), heightened apoptosis, thrombus formation, intracellular calcium accumulation, and dysregulation of neuroprotective and apoptotic gene expression were among the major impairments brought on by stroke induction. All of these findings supported the development of ischaemic damage and related neuronal stress in zebrafish larvae.

Recovery outcomes were significantly enhanced by treatment with TD:Aspirin formulations, especially at a 1:100 ratio. While morphological evaluations showed few abnormalities and better heart rate stability, behavioural analyses showed greater activity and decreased erratic swimming, indicating functional recovery. Calcium imaging demonstrated normalisation of intracellular calcium levels, indicating restoration of NMDAR function and decreased excitotoxicity, and O-dianisidine staining confirmed decreased thrombus development. At the molecular level, TD:Aspirin altered the production of neurotrophic factors, such as BDNF and GDNF, to promote neuronal survival and regeneration and successfully suppressed ROS generation and apoptosis, as seen by decreased expression of caspase 3 and caspase 9. Additionally, normalised α-tubulin and MBP expression demonstrated better cytoskeletal stability and myelination, whereas modulation of GRIN2B and SLC8A showed restoration of excitatory signalling and calcium homeostasis.

Because of the DNA nanocages’ improved bioavailability, prolonged release, and targeted distribution, TD:Aspirin had better neuroprotective effects than free aspirin and TD-only therapies. Higher drug loading maximises therapeutic efficacy without causing toxicity, as the 1:100 ratio showed especially strong results. MD simulation provides clear evidence that Aspirin intrudes into the DNA grooves by replacing water molecules and binding with TD in multimode (e.g., major groove, minor groove, backbones, outer corner mode, intercalation).

All these results point to TD: Aspirin as a potentially effective nanotherapeutic approach for ischaemic stroke that can lower oxidative stress, reduce apoptosis, restore neuronal function, and encourage vascular and neural recovery. This study concludes by showing that aspirin delivered via DNA nanocage can successfully reduce ischaemic injury in zebrafish larvae, providing a unique platform for targeted neuroprotection. These findings offer a solid basis for more preclinical testing and possible translational development of treatments for stroke and other neurovascular conditions based on DNA nanostructures.

## Acknowledgements

The authors would like to express their sincere gratitude to all members of the A.K. and D.B. groups for their critical evaluation of the work and insightful comments. K.K. expresses gratitude to GoI and DST for the National Postdoctoral Fellowship in Nanoscience and Technology. SM acknowledges the SRF fellowship from CSIR, India. PKM thanks the Department of Science &Technology (DST), India, for financial support and the Science and Engineering Research Board (SERB) for financial and computational support through CRG/2021/003659. D.B. acknowledges research funds from DBT-EMR, Gujcost-DST, and GSBTM; MoES for the STARS award; IITGN for the startup grant; and SERB and GoI for the Core research grant. A.K. thanks the GSBTM (GSBTM/JD(R&D)/663/2023-24/02003699) and ICMR (IIRP-2023-1078) for their financial assistance.

## Author Contributions

K.K. conceived the idea, planned all the experiments, analysed the results and wrote the original manuscript. S.C. and K.J. executed all in vivo experiments in zebrafish. S.M. executed the MD simulation. P.K.M. provided the simulation facility. A.K. provided the zebrafish facility. A.K. and D.B. contributed the facility and required funding. All the authors discussed the data and helped in writing and critically reading the manuscript.

